# Shared and niche-specific transcriptional signatures of macrophage aging revealed by a cross-tissue meta-analysis

**DOI:** 10.64898/2026.02.03.703654

**Authors:** Ella Schwab, Eyael Tewelde, Leon Chen, Bérénice A. Benayoun

## Abstract

**Background:** Aging is accompanied by widespread transcriptional remodeling across tissues, yet how aging impacts different categories of tissue-resident macrophages is not well understood. Macrophages are highly specialized innate immune cells shaped by their local microenvironments, suggesting that aging may elicit both shared and niche-specific transcriptional responses. Here, we performed a meta-analysis of publicly available bulk and single cell RNA-sequencing datasets to characterize age-associated transcriptional changes in murine macrophages across tissues and sexes. We curated and uniformly processed 33 macrophage transcriptomic datasets, derived from 10 distinct tissue niches, in male and female C57BL/6 mice, examining transcriptional changes as a function of age.

**Results:** The similarity of differentially expressed aging genes was compared across niches and pathway-level analysis uncovered conserved age-associated signatures, including upregulation of gene sets related to antigen presentation, antioxidant responses, and negative regulation of ferroptosis, alongside downregulation of gene sets related to Wnt, GTPase, and extracellular matrix organization signaling. Transcription factor activity inference identified consistent age-associated activation of AP-1 (Fos, Jun), C/EBPβ, PU.1, and Egr1 across niches. Meta-analysis defined 593 consistently age-altered genes in >3/4 of analyzed datasets, converging on dysregulation of small GTPase signaling. Focused analysis of alveolar macrophages and microglia, made possible by the larger number of available datasets, revealed sex-specific transcriptional programs altered with age in these macrophage subtypes.

**Conclusions:** These findings demonstrate that macrophage aging is shaped by both tissue niche and sex and provides a framework for understanding the transcriptomic signatures of macrophage aging across tissues.

## BACKGROUND

Aging results in widespread transcriptomic dysregulation across tissues [1, 2]. Pathways commonly dysregulated with age tend to correspond with canonical “hallmarks of aging”, including changes to inflammatory genes, mitochondrial dysfunction, and loss of proteostasis [3, 4]. Although some age-associated transcriptomic changes are global, many are largely tissue-specific [2, 5–7]. Both the cellular composition and the gene expression profiles of different cell types within any given tissue vary with age [8]. For example, in aged mouse microglia, both MHC class I genes and interferon-responsive genes are upregulated, which then contributes to an increase in the population of pro-inflammatory microglia in the brain with age [8, 9]. The rate at which cells and tissues age at the transcriptomic level is also asynchronous, with specific cells/tissues aging at different rates [10]. Notably, the spleen undergoes substantial transcriptomic changes earlier in life compared with other tissues [4].

Macrophages are a key innate immune cell type, responsible for various key functions such as phagocytosis, cytokine production, and antigen presentation [11, 12]. Importantly, there are two broad ontogenies for macrophages: (i) tissue-resident macrophages, which are differentiated during embryogenesis, and (ii) monocyte-derived macrophages, which differentiate throughout life in response to immunological needs [13, 14]. Although both macrophage types can cover all key functions, tissue-resident macrophages tend to play a crucial role in tissue homeostasis, growth, regeneration, and host defense through immune surveillance [13, 15]. Most adult tissue-resident macrophages, including Kupffer cells, microglia, Langerhans cells, adipose tissue macrophages, and alveolar macrophages, originate from fetal yolk sac–derived erythro-myeloid progenitors. Unlike monocyte-derived macrophages, tissue resident macrophages are long-lived and maintained through local self-renewal [14, 15]. They exhibit niche specificity, perform organ-specific functions, and are shaped by their resident tissue through local transcriptional and epigenetic programs [16, 17]. This specificity suggests that tissue-resident macrophages may undergo distinct age-associated changes, potentially leading to impaired regulation of local immune responses in aged tissues.

With age, there is a general decline in immune function known as immunosenescence. Aging results in distinct changes in the immune system that impair its capacity to respond to pathogens, maintain tissue homeostasis, and resolve inflammation [18]. Macrophages, as key innate immune effectors, are central players in this process due to their roles in antigen presentation, cytokine production, and clearance of senescent cells. Generally, in murine macrophages, pathways related to phagocytic activity, cytokine production [19], antibacterial defense [20, 21], and wound repair [22] are altered with age [23]. Understanding how macrophages function and how their transcriptional profiles shift with age is essential to dissecting their contributions to inflammaging and age-related diseases. These cells can influence the progression of several age-related diseases, including neurodegeneration, osteoporosis, atherosclerosis, cancer, diabetes, and arthritis [12, 24]

Few studies have described macrophage aging transcriptomic signatures across tissue niches [16, 25], and most meta-analyses tend to focus on age-related changes in just one specific niche. A wealth of transcriptomic data on aging macrophages has been generated by the research community and is publicly available in genomic databases [22, 26–49]. Tissue-resident macrophages perform diverse biological roles depending on their tissue context and are not a uniform population. However, it remains unclear how macrophages from different tissues change at the transcriptomic level with age, whether there are common principles to macrophage aging, or if each niche environment may shape distinct, specific aging trajectories. Thus, it is important to characterize niche-specific transcriptomic changes across both sexes and with age.

Here, we searched publicly available RNA-seq datasets of purified mouse macrophages with aging to perform a meta-analysis study of macrophage aging transcriptomic signatures using mouse as a model. Through our database search, we identified 29 mouse macrophage RNA-seq datasets, derived from 10 different niches, to be used for meta-analysis for investigation of shared *vs.* niche-specific gene regulation changes with aging (**Supplementary Table S1A**). We obtained these datasets and subjected them to standardized processing and quality control filtering. After stringent quality control and filtering (**Fig. 1A**; see Methods), we focused our analysis on 24 high-quality mouse transcriptomic datasets of purified macrophages across 10 niches – segregating datasets as a function of sex, since it impacts macrophage transcriptomes [27] (**Fig. 1A**; **Supplementary Table S1A**; see Methods).

**Figure 1.**
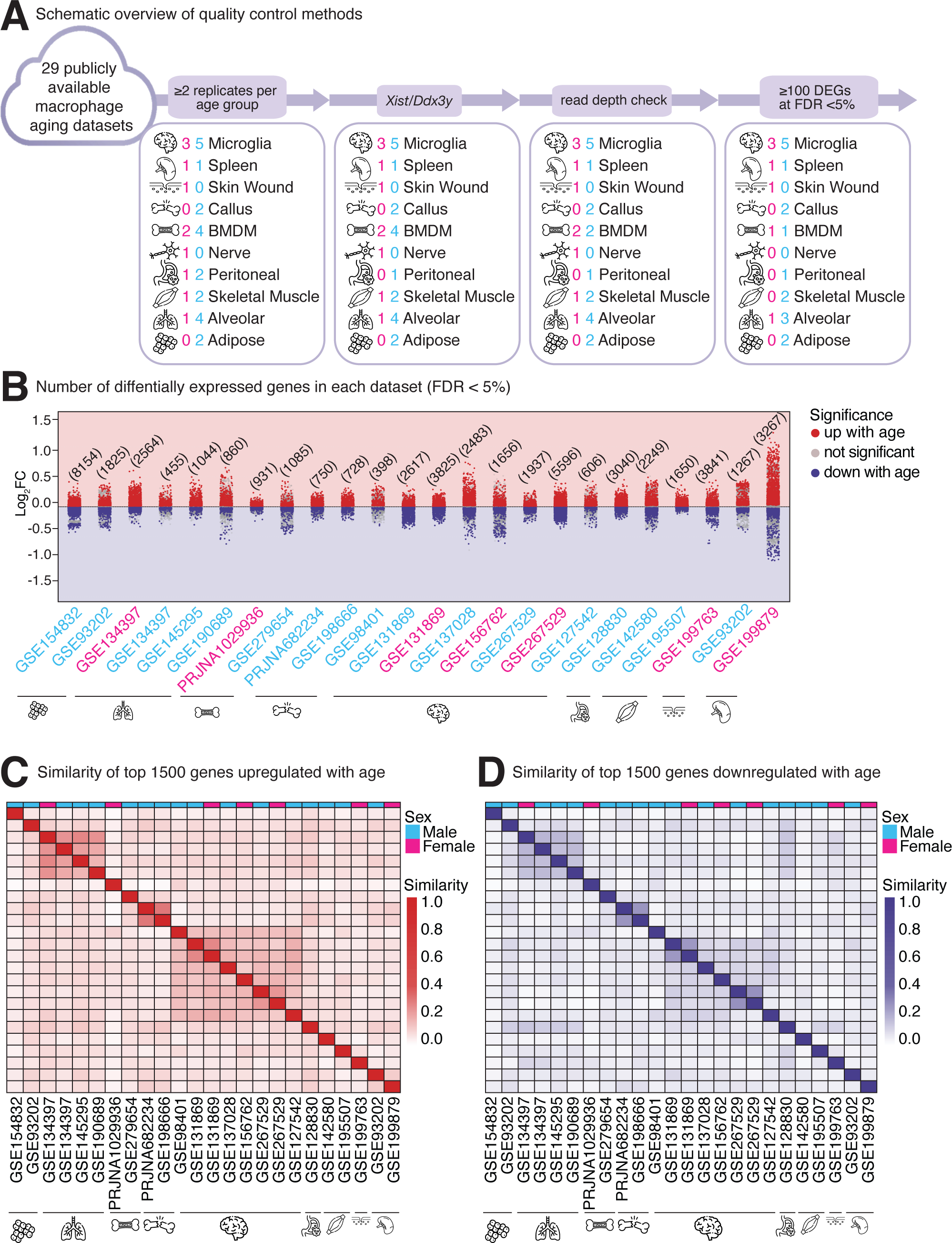
Dataset quality control and analysis of aging-associated differential gene expression across macrophage niches. **(A)** Schematic overview of data acquisition and quality control by filtering for number of replicates, sample sex, read depth discrepancy, and number of differentially expressed genes. **(B)** Jitter plot showing the total number of differentially expressed genes per macrophage transcriptomic dataset at FDR < 5%. Log_2_FC values for DEGs represent an increase (+) or decrease (-) in gene expression with a one-month increase in mouse age. Genes are colored red to indicate significant upregulation with age, blue for significant downregulation with age, and grey for genes that are not significantly altered with age. **(C)** Jaccard index heatmap displaying similarity scores of the 1500 most upregulated genes with age in each dataset, regardless of significance threshold. **(D)** Jaccard index heatmap displaying similarity scores of the 1500 most downregulated genes with age in each dataset, regardless of significance threshold.

## RESULTS

### Aging elicits transcriptional changes across murine macrophage populations of male and female mice

Our search across repositories initially identified 29 publicly available transcriptomic datasets of mouse macrophages with aging (**Supplementary Table S1**), using as criteria that: (i) the transcriptomic profiling methodology be bulk or single cell RNA-seq, (ii) the cells must be either experimentally purified (bulk RNA-seq) or purifiable *in silico* (single-cell RNA-seq), and (iii) there should be a clear age spread between groups to evaluate the effect of aging. We note that, since datasets were curated from public databases, age groups are quite variable with “young” animals ages ranging from 2 to 6 months and “old” animals ages ranging from 10 to 26 months (**Supplementary Table S1A**). Importantly, all identified datasets used variants of the C57BL/6 mouse genetic background (**Supplementary Table S1B**), which is the most standardly used genetic background to study aging in mice. Identified datasets were comprised of 6 bone marrow-derived macrophage datasets [26–29], 5 microglia datasets, [28, 30–35], 5 alveolar macrophage datasets [36–39], 3 skeletal muscle macrophage datasets [40, 41], 2 adipose macrophages datasets [42, 43], 2 peritoneal macrophage datasets [44, 45], 2 spleen macrophage datasets [42, 46], 2 bone callus macrophage datasets [47, 48], 1 skin macrophage dataset [22], and 1 nerve-associated macrophage dataset [49] (**Supplementary Table S1A**). After splitting datasets according to the author-reported biological sex of samples, so as to avoid confounding signal from potential sex-specific transcriptomic regulation [27], we started with 33 macrophage aging transcriptomic datasets for our meta-analysis.

These datasets were then processed using a standardized pipeline (see Methods), with mapping and expression quantification at the gene level using mouse genome reference mm10 (see Methods). We established a set of stringent quality filters to retain only datasets with highest confidence: (i) minimum number of replicates per age group (N ≥ 2), (ii) X– (*Xist*) and Y-linked (*Ddx3y*) sex gene markers consistency (*i.e*. agreement between transcriptome-inferred sex and author-reported sex, and consistency of sex between young and old samples) **(Supplementary Figure 1A)**, (iii) gene-mapped read depth consistency (comparable normalized read depth between the oldest and youngest age groups in each dataset), and (iv) the presence of ≥100 significantly differentially expressed genes at FDR < 5% (to guarantee there is sufficient signal *vs*. noise to process in the data). After these filters, 9 datasets were eliminated from further analysis (**Figure 1A; Supplementary Table S1**), leaving us with a total of 17 high-quality datasets from male mice and 7 high-quality datasets from female mice, for a total of 24 datasets.

Interestingly, we observed a wide range of significant differentially expressed genes (DEGs) across datasets (398 to 8154 genes at FDR <5%; **Figure 1B; Supplementary Table S2**), likely due to a range of factors both technical (*e.g*. replicate number, sequencing depth, batch effects, etc.) and biological (*e.g.* aging, niche, sex, etc.) which influence discovery power. Importantly, we used a linear modeling approach for age expressed in months in our analysis so that (i) datasets with 3 or more age groups could use the full statistical power of such experimental designs, and (ii) reported log_2_(fold-change) values would be expressed by unit of time and thus directly comparable across datasets (**Figure 1B**). To note, various datasets showed varying ranges of log_2_(fold-change per month) values, suggesting that macrophage transcriptomes of various niches may have different sensitivity to aging – potentially as a function of sex. Alternatively, differences in covered age ranges (see above) may differentially impact estimates, as the rate of change in gene expression may not be constant across the lifespan [50] [51].

To assess the similarity of age-related transcriptomic changes across niches, we identified the top 1,500 genes showing the strongest age-associated upregulation and downregulation, ranked by log_2_(fold-change per month), regardless of adjusted p-value (**Supplementary Table S3**), to have the fairest possible pairwise comparison of datasets. We then calculated Jaccard index similarity scores separately for upregulated and downregulated genes in each dataset to determine the degree of sharing of transcriptional responses across macrophage niches (**Figure 1C,D**). Importantly, our analysis revealed that alveolar macrophage datasets and microglia datasets exhibited the highest similarity scores for both genes upregulated **(Figure 1C)** and downregulated **(Figure 1D)** with age, potentially reflecting the greater statistical power afforded by the larger number of datasets available for these niches. Interestingly, within the two niches, upregulated genes showed greater cross-dataset similarity than downregulated genes, potentially suggesting that the transcriptional response to aging is more constrained and conserved in its activation programs than in its repression programs. Our observations thus show that: (i) as expected, macrophages within any specific niche age most similarly to macrophages from the same niche, regardless of sex or specific age range, and (ii) there is only modest overlap in age-regulated genes between macrophages of different niches, suggesting that aging impacts each niche in specific, divergent ways. Importantly, even with a more restrictive definition, using Jaccard indices for all age-upregulated or downregulated genes at FDR < 5% we observed largely consistent results **(Supplementary Figure 2A,B**). Thus, our analyses suggest that macrophage transcriptional remodeling in response to aging is largely niche-specific, at least at the gene-level.

### Multi-contrast enrichment analysis reveals pan-macrophage aging transcriptional response signatures across niches

Since similar functional outcomes can be achieved by regulating different sets of genes within the same pathway, we next asked whether macrophage aging could lead to the regulation of similar pathways across niches, even in the context of limited gene-level sharing. Specifically, we leveraged a multi-contrast functional enrichment paradigm, mitch [52], whereby each dataset was used as its own contrast in the analysis, to identify significantly enriched gene sets across our meta-analysis datasets **(Figure 2A, 3A; Supplementary Figure S3; Supplementary Table S4)**. Importantly, we focused on significant top gene sets and pathways with largely consistent directionality across datasets, using high-quality, well-characterized gene sets from Gene Ontology, Reactome, and KEGG (see Methods). We reasoned that this approach would help identify core pathways regulated across niches in response to aging in murine macrophages.

**Figure 2.**
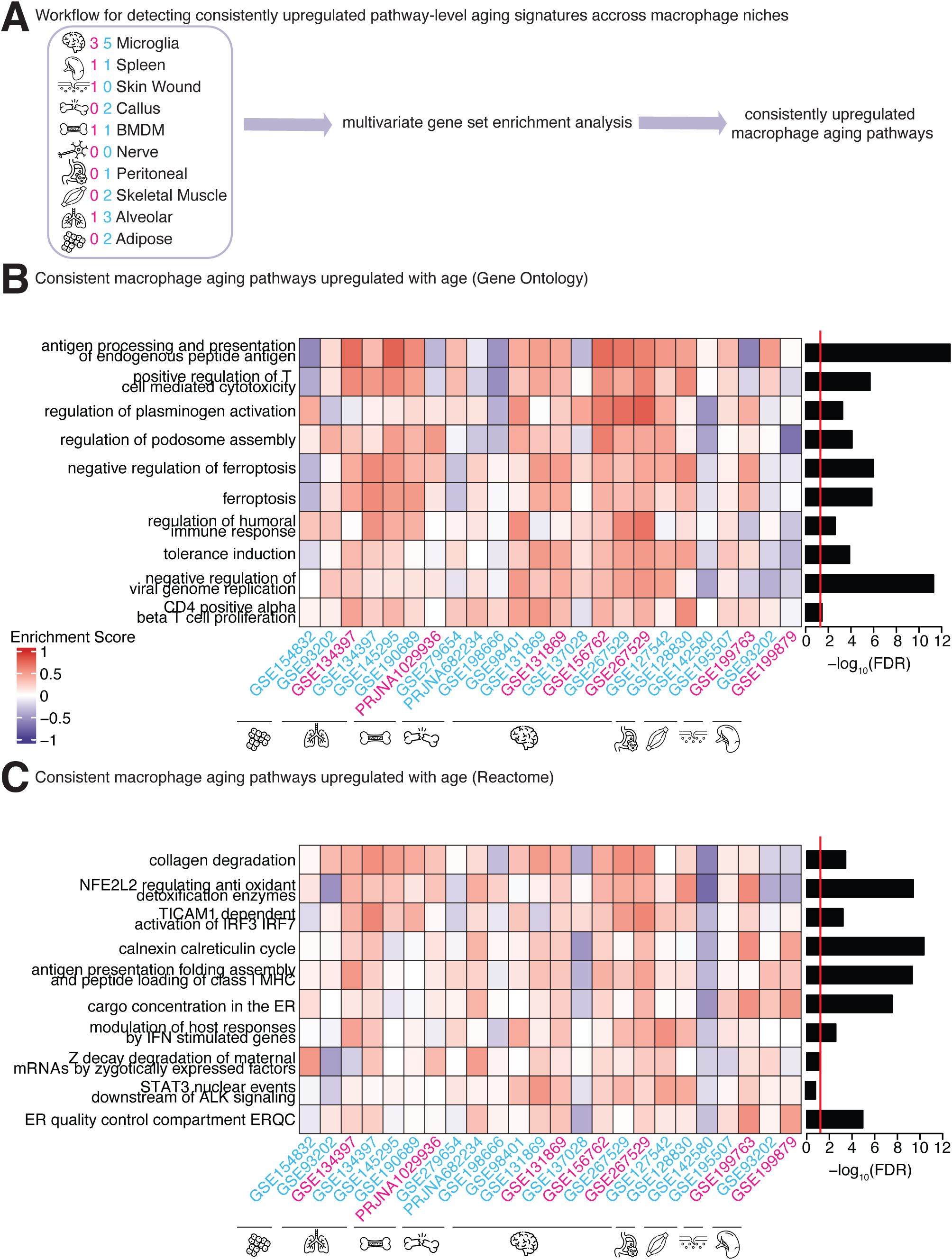
Multivariate Gene Set Enrichment Analysis of age-associated transcriptomic changes across datasets for pathways upregulated with age in macrophages. **(A)** Diagram showing the workflow for detecting commonly upregulated aging pathways. Pathways were classified as upregulated (red) when the direction of enrichment (enrichment score) is consistent in at least 18 out of 24 datasets. **(B)** Heatmap for upregulated Gene Ontology Biological Process (GOBP) pathways with age across macrophage niches at FDR < 5%. **(C)** Heatmap for upregulated Reactome pathways with age across macrophage niches. The red line indicates FDR < 5%.

Interestingly, gene sets and pathways related to antigen processing, regulation of plasminogen activation, and regulation of humoral immune response were upregulated across datasets **(Figure 2B,C)**. In addition, ferroptosis-related terms were increased in most datasets, as well as antigen presentation. To note, negative regulation of ferroptosis was exceptionally upregulated in microglia and alveolar macrophages **(Figure 2B).** Collagen degradation and *Nfe2l2*-regulating antioxidant detoxification enzymes were upregulated in nearly every niche except spleen **(Figure 2C)**. Ferroptosis has been increasingly linked to aging through age-associated iron accumulation and oxidative stress, both of which can promote ROS production and inflammatory signaling [53, 54]. In macrophages, elevated ROS may contribute to inflammaging, necessitating adaptive responses to limit ferroptosis. The upregulation of negative regulators of ferroptosis, particularly in microglia and alveolar macrophages, together with increased *Nfe2l2*-driven antioxidant pathways, suggests a compensatory mechanism to counteract oxidative damage during aging. In parallel, widespread upregulation of collagen degradation pathways indicates enhanced extracellular matrix remodeling, consistent with chronic inflammatory or stress-responsive tissue environments.

Next, we explored gene sets and pathways with consistent age-related downregulation **(Figure 3; Supplementary Table S4)**. Interestingly, gene sets related to positive regulation of extracellular matrix organization, negative regulation of *Flt3* signaling, and GTPase signaling were downregulated with age across most macrophage niches **(Figure 3B,C)**. The downregulation of negative regulators of Flt3 signaling is consistent with a net increase in Flt3 signaling activity with age. Flt3 signaling is characteristic of monocyte-derived macrophage lineages but absent from embryonic-derived tissue-resident alveolar macrophages. Longitudinal fate-mapping confirms that HSC-derived cells progressively contribute to the alveolar macrophage niche in an age-dependent manner [55]. Thus, age-related downregulation of *Flt3* signaling in bulk macrophage transcriptomes may reflect diminished recruitment of new monocyte-derived macrophages during aging in niches where tissue resident macrophage pools are sufficiently able to maintain their population or self-renew. Thus, the age-related downregulation of negative *Flt3* regulators in bulk macrophage transcriptomes may reflect increased recruitment of monocyte-derived macrophages with age, consistent with the progressive replacement of tissue-resident macrophage pools by HSC-derived cells across niches. Several types of Rho GTPases are expressed within the myeloid lineage including RhoQ, Rac1, Rac2, RhoA and RhoC. These GTPases are important for regulation of actin cytoskeleton dynamics and thus cell adhesion, and migration [56]. Although the role of GTPase signaling in macrophage aging is not well-defined, aging is known to disrupt the integrity of the extracellular matrix [57, 58], and the downregulation of GTPase-related gene sets may reflect broader cytoskeletal and membrane trafficking changes associated with macrophage functional decline with age.

**Figure 3.**
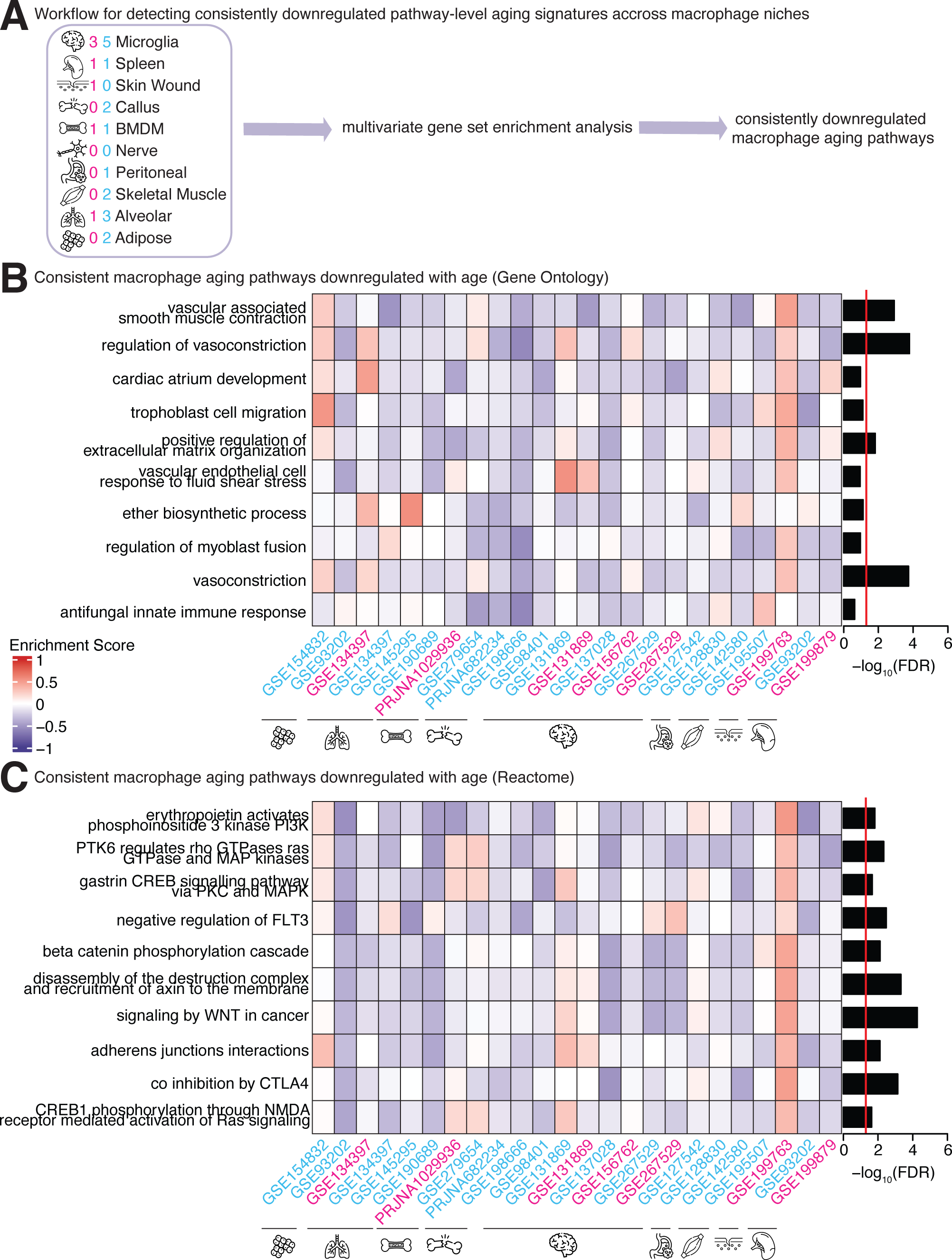
Multivariate Gene Set Enrichment Analysis of age-associated transcriptomic changes across datasets for pathways downregulated with age in macrophages. **(A)** Diagram showing the workflow for detecting commonly downregulated aging pathways. Pathways were classified as downregulated (blue) when the direction of enrichment (enrichment score) is consistent in at least 18 out of 24 datasets. **(B)** Heatmap for downregulated Gene Ontology Biological Process (GOBP) pathways with age across macrophage niches at FDR < 5%. **(C)** Heatmap for downregulated Reactome pathways with age across macrophage niches. The red line indicates FDR < 5%.

Wnt signaling was downregulated in the Reactome and KEGG databases, supporting a consistent downregulation of Wnt signaling across macrophage niches **(Figure 3B; Supplementary Figure 3B)**. Wnt signaling is known to regulate macrophage polarization [59], so this loss of Wnt signaling during aging may contribute to the maladaptive shift towards pro-inflammatory M1 polarization that occurs with age [60, 61].

Together, our global functional enrichment analysis highlights the existence of modest but consistent patterns of age-associated transcriptional changes across macrophage niches. These results show both conserved and niche-specific features of macrophage aging and will help deepen our understanding how specific signaling pathways may contribute to age-related changes in murine innate immunity, regardless of niche.

### Specific transcription factors show altered activity across murine macrophage niches with age

Next, we asked whether we could pinpoint master regulators explaining transcriptional aging of macrophages across niches **(Figure 4A)**. To infer which transcription factors (TFs) might regulate these pan-macrophage aging genes, we performed TF activity analysis using the decoupleR algorithm [62] to identify the top TFs with predicted activity change with aging in ≥18 out of our 24 datasets **(Figure 4A; Supplementary Table S5)**. The TFs most consistently predicted to be activated with age across datasets were Jun and Cebpb. *Jun* encodes c-Jun, a member of the Jun protein family that forms the AP-1 transcription factor, and is involved in positive regulation of the cell cycle [63, 64], cell proliferation [65, 66], and immune responses [64].

**Figure 4.**
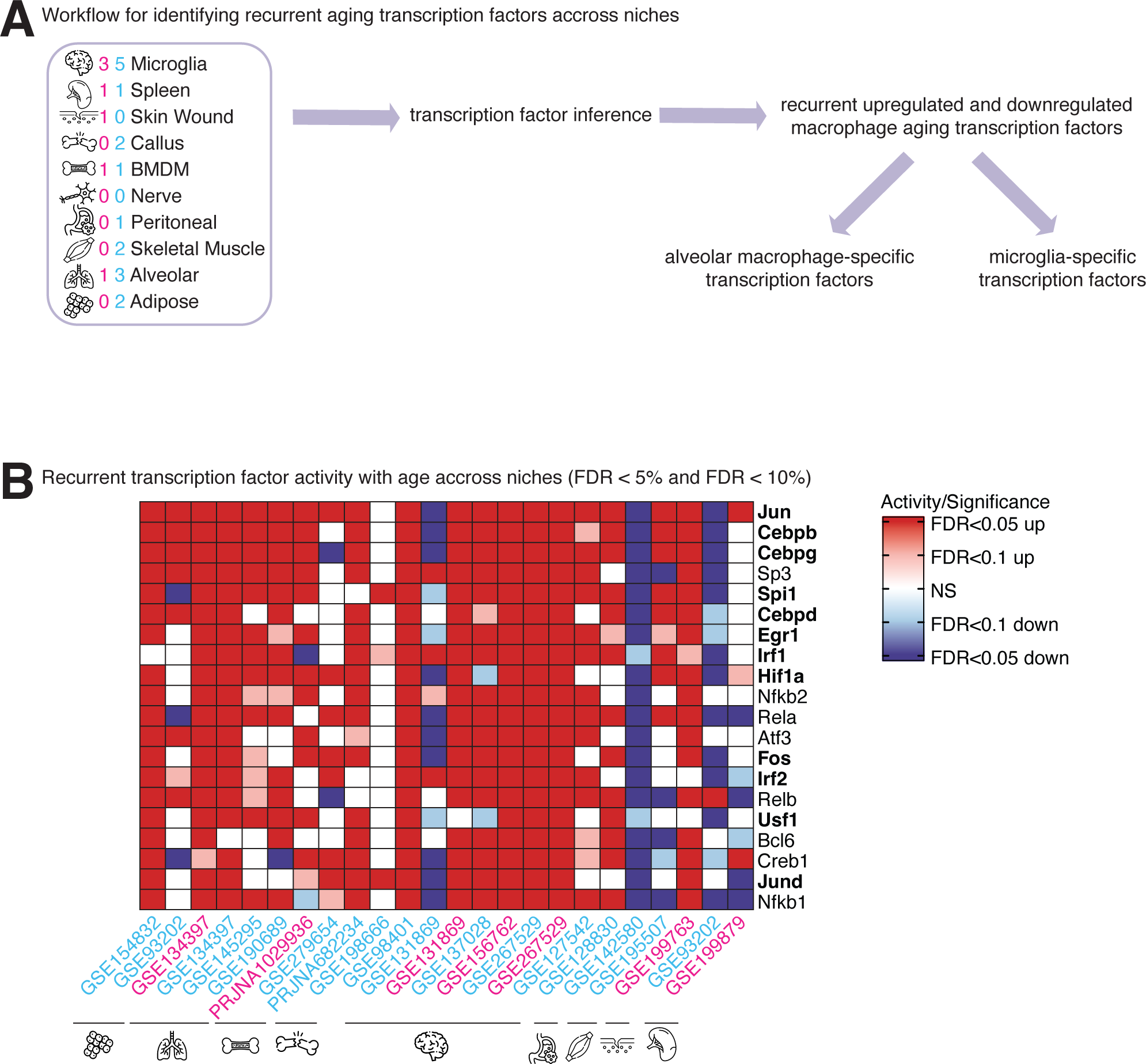
Recurrent predicted changes in transcription factor activity with aging across macrophage niches. **(A)** Schematic overview of transcription factor activity inference analysis across all macrophage niches. **(B)** Heatmap of recurrent transcriptional factor activity inferred across macrophage niches, with colors indicating significant activation or repression (red = activated, blue = repressed). The corresponding significance of activation or repression is evaluated at thresholds of FDR < 5% and < 10%, where darker tones indicate greater significance in their respective direction.

*Cebpb* encodes C/EBPβ, a transcription factor of the CCAAT/enhancer-binding protein (C/EBP) family that is expressed in macrophages [67]. Under homeostatic conditions, C/EBPβ activity is relatively low; however, its expression and transcriptional activity are induced by inflammatory cytokines, lipopolysaccharide (LPS), and interferon-γ (IFN-γ), linking C/EBPβ to innate immune activation [68] Upon activation, C/EBPβ regulates gene programs associated with macrophage polarization, including M2-associated genes, and contributes to inflammatory responses in multiple tissues [69], including the central nervous system, where it has been implicated in neuronal inflammation [70]. An increase in predicted activity of C/EBPβ with age in tissue resident macrophages may reflect a response to persistent low-grade inflammation. The predicted activity of *Spi1*, which encodes the pioneering transcription factor PU.1, also increases with age across most macrophage niches. PU.1 together with C/EBP family members establishes and maintains the majority of macrophage-specific enhancers during lineage commitment [71], enrichment of these pioneering factors in our analysis likely reflects changes in macrophage cellular identity with age, a phenomenon that has been previously reported [72]. PU.1 is essential for tissue-resident macrophage development [73], regulates both microglial maintenance [74] and inflammatory responses [72], and altered PU.1 activity has been linked to Alzheimer’s disease in humans [75].

Interestingly, the predicted activity of Egr1, a known regulator of macrophage inflammatory gene expression [76], is also consistently increased across most macrophage niches **(Figure 4B)**. In contrast, both spleen datasets generally showed downregulation of TFs that were upregulated in other macrophage niches. This pattern indicates that spleen macrophages may follow a unique transcriptional trajectory with age, potentially reflecting tissue-specific differences in the microenvironment.

### Transcription factor-pathway network analysis reveals coordinated transcriptional regulation of macrophage aging signatures

Next, to investigate whether TFs with altered activity across macrophage niches with aging converge on our identified commonly identified aging-associated pathways, we constructed a TF-gene-pathway network by calculating the overlap between transcription factor target genes and genes driving age-associated pathway enrichment **(Figure 5A, Supplementary Figure 4A;** see Methods). Across both upregulated and downregulated macrophage aging pathways, Jun targets showed the broadest overlap, suggesting that Jun-mediated transcriptional regulation may coordinate a large proportion of the age-associated gene expression changes observed across macrophage niches. In addition, Hif1a had the most target genes in common with vasoconstriction-related pathways **(Figure 5B)**, consistent with its established role in macrophage biology. HIF-1α is a regulator of macrophage immune responses, migration, and metabolic reprogramming [77, 78], and its activity is induced under both hypoxic and inflammatory conditions through NF-κB-dependent transcription and mTOR signaling. Given that aging is associated with chronic low-grade inflammation and tissue hypoxia, the enrichment of Hif1a target genes in vasoconstriction-related pathways may reflect an age-associated shift toward pro-inflammatory macrophage states, consistent with HIF-1α’s known role in promoting M1 polarization through glycolytic reprogramming [79]. Creb1 targets showed a higher degree of overlap with pathways related to Wnt signaling, axin recruitment, and β-catenin phosphorylation **(Figure 5C)**. In macrophages, CREB1 promotes anti-inflammatory M2 polarization through a CREB, C/EBPβ transcriptional cascade, whereby CREB-mediated induction of *Cebpb* is required for the coordinated upregulation of M2-specific genes, while leaving M1 inflammatory gene expression intact [69]. The CREB/CRTC pathway has similarly been shown to drive M2 polarization downstream of prostaglandin E2 signaling, with macrophage-specific disruption of CREB reducing M2 marker expression [80]. The enrichment of Creb1 targets in downregulated Wnt-related pathways is therefore notable, as reduced Wnt-CREB signaling activity with age may impair this anti-inflammatory cascade and contribute to the age-associated shift away from M2 macrophage polarization states observed across niches. These findings suggest that the TFs predicted to drive macrophage transcriptional aging programs do not act independently, but rather converge on shared gene programs that are responsible for the functional remodeling of macrophages with age.

**Figure 5.**
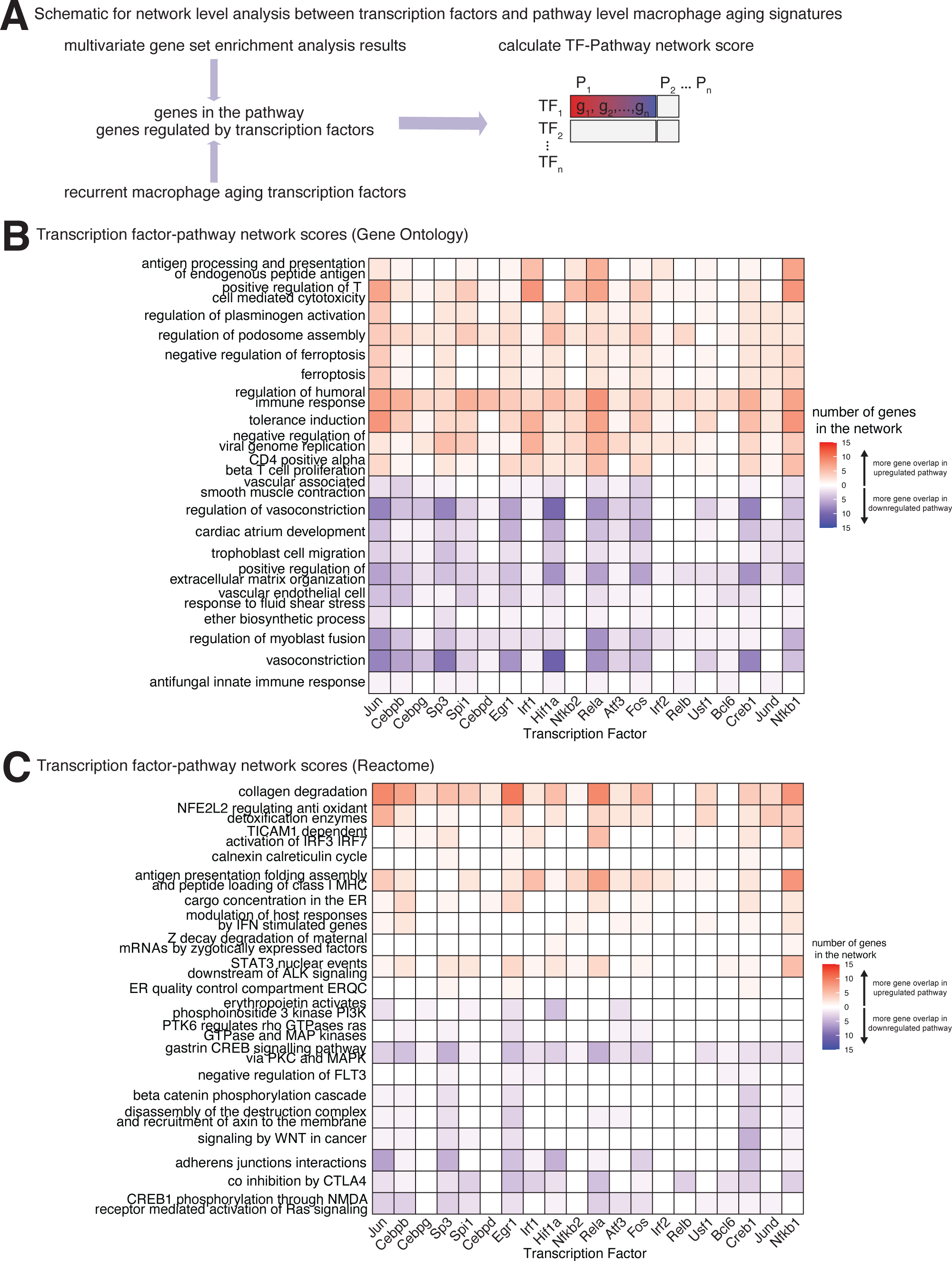
Transcription factor-gene-pathway network for common upregulated and downregulated macrophage aging signatures. **(A)** Schematic overview of network level analysis between enriched transcription factors and pathway level macrophage aging signatures. **(B)** Heatmap showing the number of genes in the transcription factor-pathway network for upregulated and downregulated Gene Ontology Biological Process (GOBP) pathways with age across macrophage niches. **(C)** Heatmap showing the number of genes in the transcription factor-pathway network for upregulated and downregulated Reactome pathways with age across macrophage niches.

### Meta-analysis reveals a core set of murine macrophage aging genes

Finally, to identify a core, pan-macrophage aging signature, we leveraged a meta-analysis approach across our murine macrophage aging datasets to identify genes consistently altered with age using the ‘metaRNAseq’ paradigm [81] (see Methods) **(Figure 6A).** Both Fisher’s combined probability test and the inverse normal method were used to define meta-level statistical significance. Across the 24 datasets, we identified 593 genes that were both statistically significant and consistently regulated in the same direction in ≥ 18 out of 24 datasets (*i.e*. 3/4 of the datasets; **Figure 6B)**. To further characterize the consistently altered macrophage aging genes and investigate their potential functional relationships, we performed enrichment analysis with Gene Ontology **(Figure 6B)**, Reactome **(Figure 6C)** and KEGG **(Supplementary Figure 5C)**. Among the 342 genes consistently upregulated with age, we found significant enrichment for terms related to extracellular space, MHC Class I antigen presentation, immune effector processes and the Golgi membrane **(Figure 6C)**. The 251 genes consistently downregulated with aging across niches showed enrichment for terms related to GTPase signaling such as RHO GTPase cycle and RAC1 GTPase cycle. Interestingly genes related to the regulation of actin dynamics were also downregulated with aging **(Figure 6D, Supplementary Figure 5C)**. Actin polymerization is known to be disrupted in alveolar macrophages [82] and with aging in general [83]. These processes are functionally interconnected, as Rho family GTPases coordinate actin cytoskeletal remodeling to support both phagocytic uptake and the vesicle trafficking steps required for cytokine secretion and antigen presentation [84]. The coordinated downregulation of GTPase regulatory activity and actin dynamics genes with age, alongside upregulation of Golgi membrane and antigen presentation terms across niches, may therefore reflect a broad age-associated remodeling of macrophage secretory and effector machinery.

**Figure 6.**
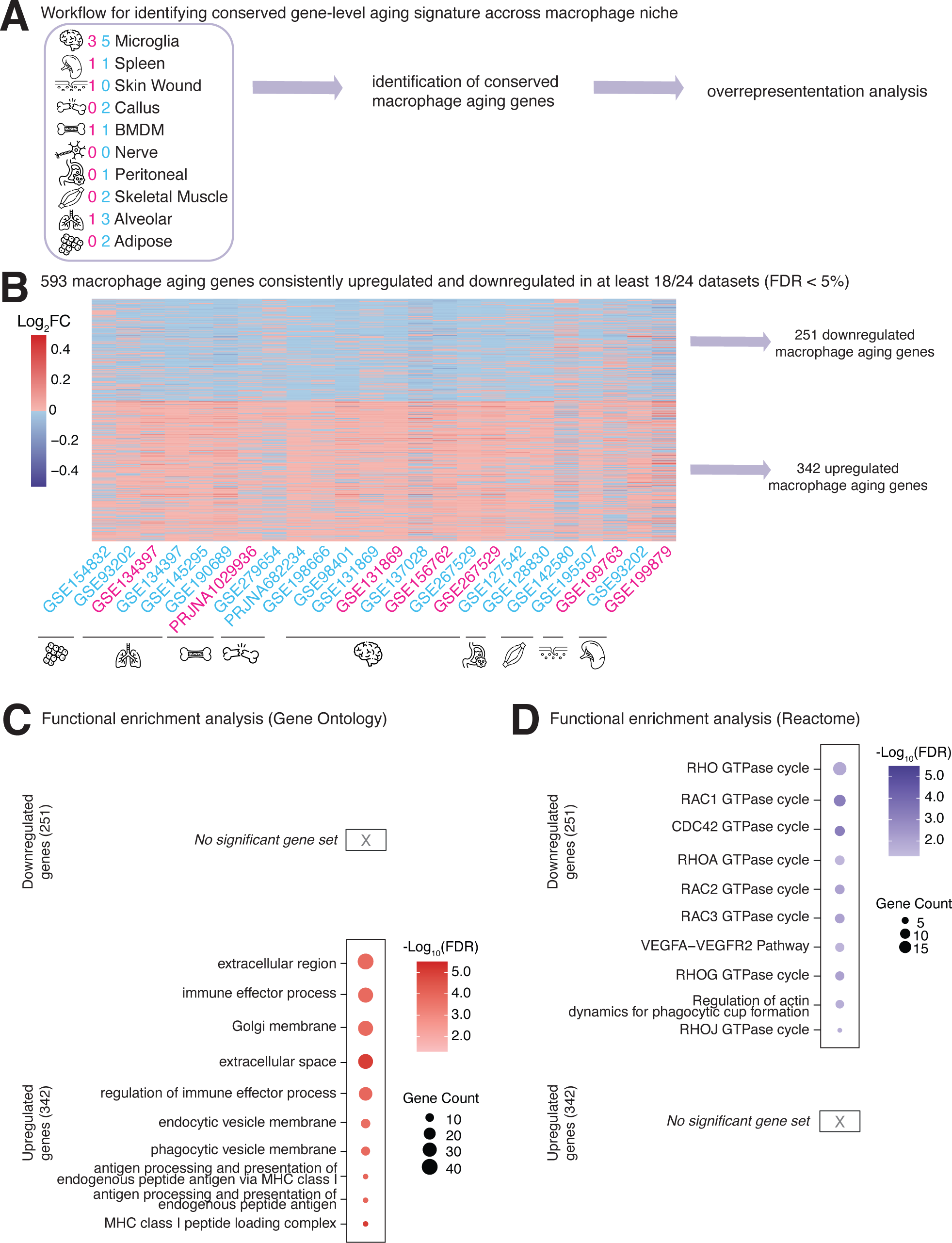
Integrated analysis of shared macrophage gene signatures across tissue niches with age, their functional convergence, and their transcriptional regulation. **(A)** Schematic overview of the pan-macrophage aging gene analysis workflow, including the identification of common aging-associated genes across macrophage niches. **(B)** Gene expression heatmap displaying a core set of age-associated macrophage genes consistent in at least 18/24 datasets across all macrophage niches, colored by log_2_ fold change (log_₂_FC) with age. Positive log_2_ fold change (log_2_FC) values indicate increased expression with age, and negative log_2_FC values indicate decreased expression with age. **(C)** Gene ontology enrichment analysis of the core macrophage aging gene signatures and their functional associations. **(D)** Reactome enrichment analysis of the core macrophage aging gene signatures and their functional associations.

Applying a more stringent criteria, we identified 35 genes that were both statistically significant and consistently regulated in the same direction in ≥ 21 out of 24 datasets **(Supplementary Figure 5A,B)**. There were 12 genes were consistently downregulated with age, including *Ap2a2, Arap3, Arhgap18, Caprin2, Frat2, Galnt10, Mapk14, Nrros, Plcl2, Sipa1, Tm6sf1*, and *Wdr41*. The other 23 significant meta-level genes regulated with aging across murine macrophages were consistently upregulated **(Supplementary Figure 5B**). Among these highly consistent macrophage aging genes (*e.g*. *Mapk14, Csf1, Nrros, Il18bp)*, many have known roles in macrophage biology, inflammation, or aging generally [85–89]. First, among the genes downregulated across niches, *Mapk14* or p38α is a member of the MAPK family and is known to be a regulator of inflammatory response in many different cell types. Normally, p38α is thought to promote inflammation by helping make pro-inflammatory cytokines such as TNF-α and IL-1β [90]. However, myeloid expression of p38α may also have a protective or anti-inflammatory role [91]. *Nrros* encodes a leucine-rich repeat protein expressed by microglia and other CNS myeloid cells, where it acts as a regulator of inflammatory and oxidative processes [92]. Its expression is generally downregulated across macrophage niches suggesting that the ability of tissue resident macrophages to respond to oxidative stress and regulate TGF-B1 signaling may be diminished with age [93]. *Csf1*, which is consistently upregulated with age across niches, is required for macrophage growth and survival [94], and its expression is known to change with age in mice and humans [87, 95]. Similarly, *Il18bp*, encoding IL-18 binding protein, is consistently upregulated across macrophage niches with age and is responsible for neutralizing the pro-inflammatory activity of IL-18 [96]. IL-18 levels are associated with both healthy aging and frailty index in centenarians [97] so, the higher expression of *IL18bp* might be associated with a compensatory mechanism against the IL-18 mediated pro-inflammatory signaling that occurs with age.

Several of the 35 highly consistent macrophage aging genes *Arhgap18, Nucb1, Rabif, Wdr41, Sipa1*, and *Arap3* were functionally linked through GTPase regulatory activity, indicating coordinated modulation of small GTPase signaling during macrophage aging. *Arhgap18* encodes a GTPase-activating protein (GAP) that modulates RhoA activity to regulate cell morphology, spreading, and migration [98]. *Nucb1* encodes a protein that acts as a calcium-regulated guanine nucleotide dissociation inhibitor for Gαi subunits, which are the GTPase components of heterotrimeric G protein complexes [99]. *Rabif* (Rab Interacting Factor) is a protein that regulates Rab GTPases, which control intracellular vesicle transport. *Wdr41* acts as a scaffold within the CSW complex (C9orf72–SMCR8–WDR41). The CSW complex exhibits GEF and GAP activity toward specific small GTPases, including members of the ARF and RAB families, which are important for membrane trafficking and autophagy [100]. *Sipa1*, or signal-induced proliferation-associated gene 1, encodes a GTPase-activating protein (GAP) specific for *Rap1*, functioning to downregulate *Rap1* signaling pathways that govern cell proliferation, adhesion, and survival [101]. Finally, *Arap3* (ArfGAP with RhoGAP domain, ankyrin repeat, and PH domain 3) encodes a multi-domain signaling protein that functions as a GTPase-activating protein (GAP) for both *Arf6* and Rho family GTPases. It acts as a downstream effector of PI3K signaling, linking phosphoinositide metabolism to cytoskeletal and membrane dynamics [102].

Together, these results define a core set of murine macrophage aging genes that are consistently altered across tissue niches and across sexes. This analysis identifies regulation of GTPase signaling as a potential mechanism underlying age-related changes in murine macrophage function.

### Over-representation analysis shows that alveolar macrophages and microglia exhibit distinct sex-specific transcriptional programs during aging

We next asked whether we could identify niche-specific pathways and gene sets regulated with aging, since macrophages from the same niche showed broadly colinear aging responses (**Figure 1A,C,D**). We took advantage of the existence of multiple datasets profiling macrophage aging in the brain, lung, spleen, bone marrow, bone callus, skeletal muscle and adipose tissue niches, encompassing both sexes, to identify sex-divergent and sex-convergent regulation of transcriptional aging signatures in each niche (**Figure 7, 8; Supplementary Figure 6**).

**Figure 7.**
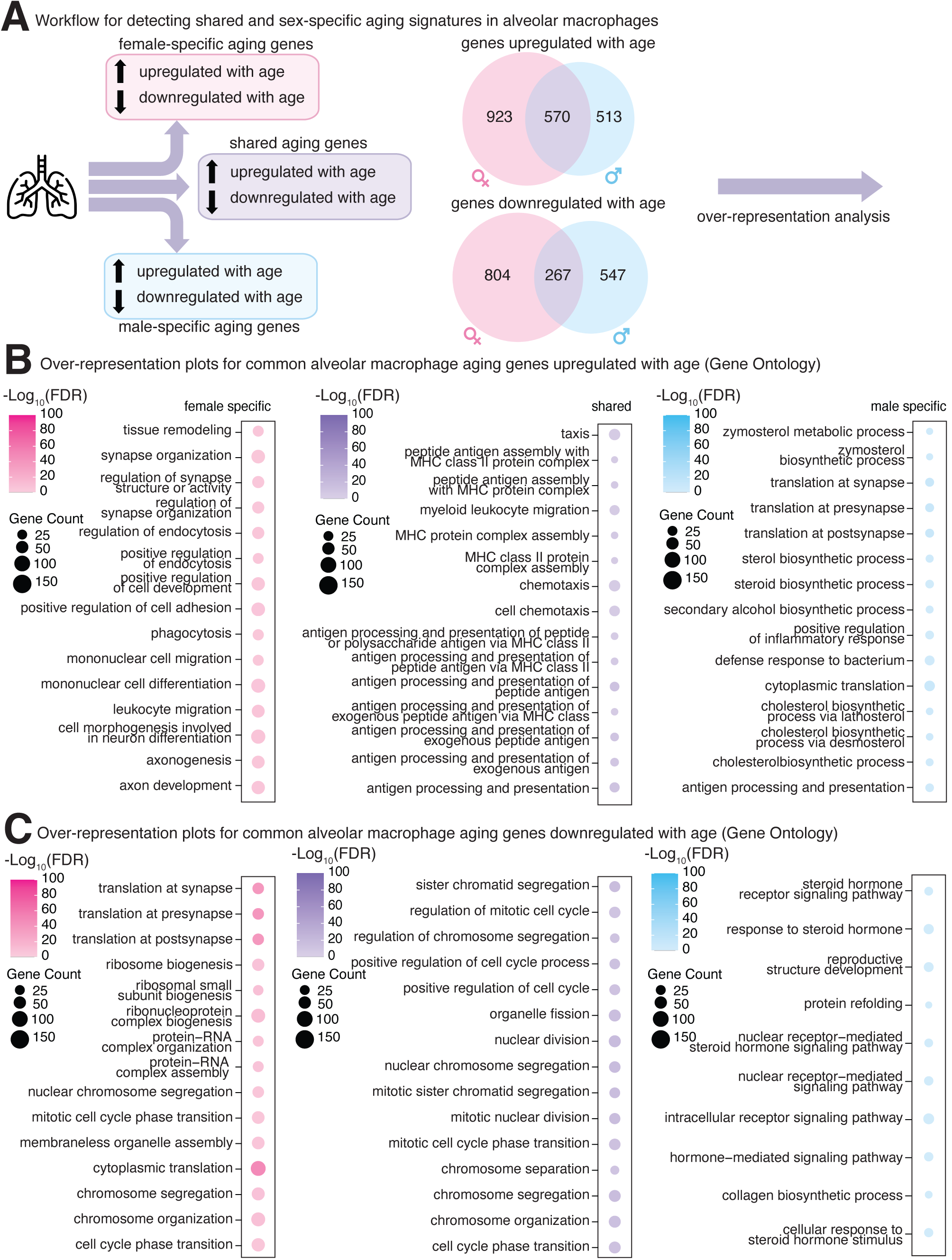
Over-Representation Analysis for differentially expressed genes with age in alveolar macrophages. **(A)** Schematic overview of Over-Representation Analysis (ORA) pipeline for differentially expressed genes in alveolar macrophages at FDR < 5% in only males, females, or both sexes. **(B)** ORA plots for differentially expressed genes at FDR < 5% that are upregulated with age in alveolar macrophages in males, females, or both sexes. **(C)** ORA plots for differentially expressed genes at FDR < 5% that are downregulated with age in alveolar macrophages in males, females, or both sexes.

**Figure 8.**
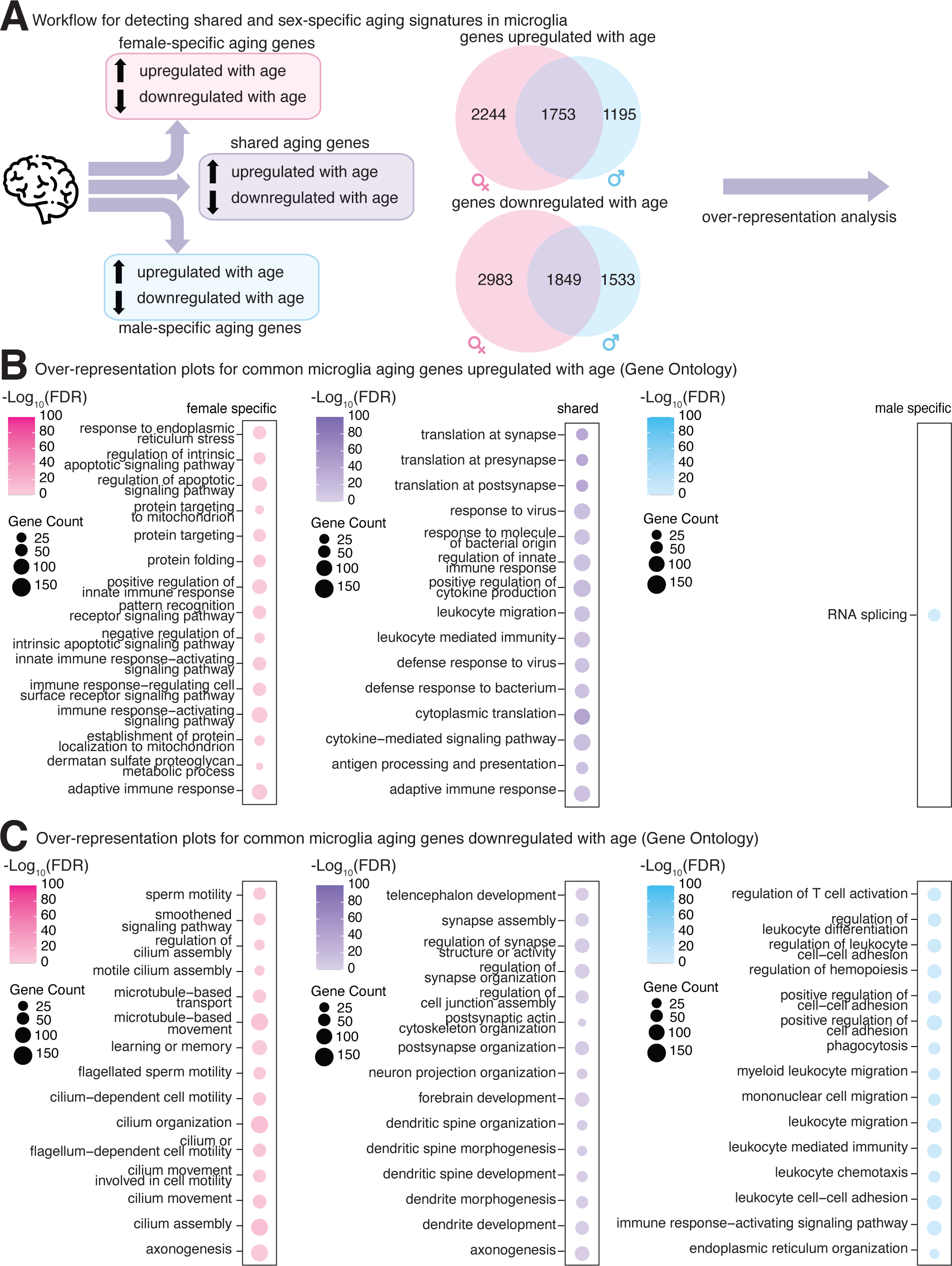
Over-Representation Analysis for differentially expressed genes with age in microglia. **(A)** Schematic overview of Over-Representation Analysis (ORA) pipeline for differentially expressed genes in microglia at FDR < 5% in only males, females, or both sexes. **(B)** ORA plots for differentially expressed genes at FDR < 5% that are upregulated with age in microglia in females only or both sexes. **(C)** ORA plots for differentially expressed genes at FDR < 5% that are downregulated with age in microglia in males, females, or both sexes.

Thanks to the large number of independent datasets, we were able to robustly identify male-specific, female-specific, and sex-shared age-regulated genes in both alveolar macrophages and microglia (**Figure 7A, 8A; Supplementary Table S7**). In contrast, there were substantially fewer shared age-regulated genes in macrophages from the other 5 niches (with only 2 datasets each; *i.e*. spleen, bone callus, skeletal muscle, bone marrow, and adipose tissue; **Supplementary Figure 6**), likely due to lower meta-analysis power. Thus, we focus on the more highly powered niches, alveolar macrophages and microglia, for further analyses below.

To better understand the functional impact of age-related gene regulation in female vs. male alveolar macrophages and microglia, we investigated the biological processes associated with commonly upregulated and downregulated genes using over-representation analysis (ORA) (see Methods). Across alveolar macrophage datasets, we identified 923 female-specific and 513 male-specific genes upregulated with age, along with 570 genes that were consistently upregulated with aging in both sexes **(Figure 7A)**. In contrast, we identified 804 female-specific and 547 male-specific genes downregulated with age, and 267 genes downregulated with aging in both sexes **(Figure 7A)**. Among alveolar macrophage datasets, genes significantly upregulated with age **(Figure 7B)** were associated with terms related to chemotaxis and antigen presentation in both sexes. In females, additional enrichment was observed for migration, phagocytosis, and endocytosis; in males, enriched terms included cytoplasmic translation, sterol biosynthesis, defense response to bacterium, and cholesterol biosynthesis. For downregulated genes **(Figure 7C)**, terms related to the cell cycle were represented in both sexes. Female datasets also showed enrichment for cytoplasmic translation, cell cycle, and ribosome biogenesis, while male datasets were enriched for steroid hormone receptor signaling, protein folding, and response to steroid hormone. Notably, pathways related to translation showed opposing age-associated regulation between sexes, being upregulated in male alveolar macrophages but downregulated in female alveolar macrophages. Together, these findings indicate sex-specific transcriptomic remodeling in aging alveolar macrophages, with female aging characterized by coordinated downregulation of translational and ribosomal pathways suggestive of impaired translational machinery. In contrast, enrichment of androgen receptor signaling in male macrophages points to a greater influence of sex hormone–dependent regulation on age-associated changes in male alveolar macrophages, rather than purely chromosomal factors.

Across microglia datasets, we identified 2,244 female-specific and 1195 male-specific genes upregulated with age, along with 1,753 genes upregulated with age in both sexes **(Figure 8A)**. In contrast, we identified 2,983 female-specific and 1,533 male-specific genes downregulated with age, and 1,849 genes downregulated with age in both sexes **(Figure 8A)**. Genes upregulated with aging in both sexes were enriched for translation, response to virus, antigen presentation, and cytokine production **(Figure 8B)**. Additionally, genes upregulated with aging in female microglia showed specific enrichment for response to endoplasmic reticulum stress, positive regulation of innate immune response, protein folding, and apoptotic signaling pathways. Genes upregulated with aging in male microglia showed enrichment for RNA splicing. Downregulated genes with age across sexes were enriched for actin cytoskeleton, synapse assembly, and dendritic spine development **(Figure 8C)**. Genes specifically downregulated in aging female microglia showed enrichment for microtubule-based movement, cilium assembly, motility, and axonogenesis, whereas those downregulated specifically in male microglia were enriched for leukocyte differentiation, migration, and T cell activation. Overall, these results demonstrate that microglial aging involves both shared and sex-specific transcriptional changes. Across sexes, aging is marked by increased immune activation and reduced cytoskeletal, synaptic, and metabolic gene expression. However, female microglia display additional enrichment of ER stress, and protein folding pathways, suggesting heightened cellular stress and proteostatic burden, whereas male microglia show preferential downregulation of leukocyte-mediated programs. These divergent patterns support the existence of sex-specific mechanisms underlying microglial functional decline during aging.

### Transcription factor inference analysis shows that alveolar macrophages and microglia exhibit distinct gene regulation programs during aging

Since we had the most power in these niches (**Figure 7, 8)**, we next asked whether alveolar macrophages and microglia showed specific TFs activity changes with aging, distinct from the niche-agnostic analysis **(Figure 9A)**. Interestingly, TFs of the E2f family (E2f1, E2f2, E2f4) showed strong predicted decreased activity in aging alveolar macrophages, whereas Cebp TFs (Cebpb, Cebpg) showed clear predicted increased activity with aging **(Figure 9B)**. Traditionally, E2f transcription factors are known to regulate cell cycle progression and cell proliferation [103]. E2f1-E2f3 are associated with G1/S transition and so this downregulation may reflect reduced proliferation and DNA synthesis with age in macrophages [103]. In microglia, several members of the Atf, Irf, and Cebp families showed predicted increased activity with age including Atf3, Cebpg, Irf1 and Irf2 **(Figure 9C).**

**Figure 9.**
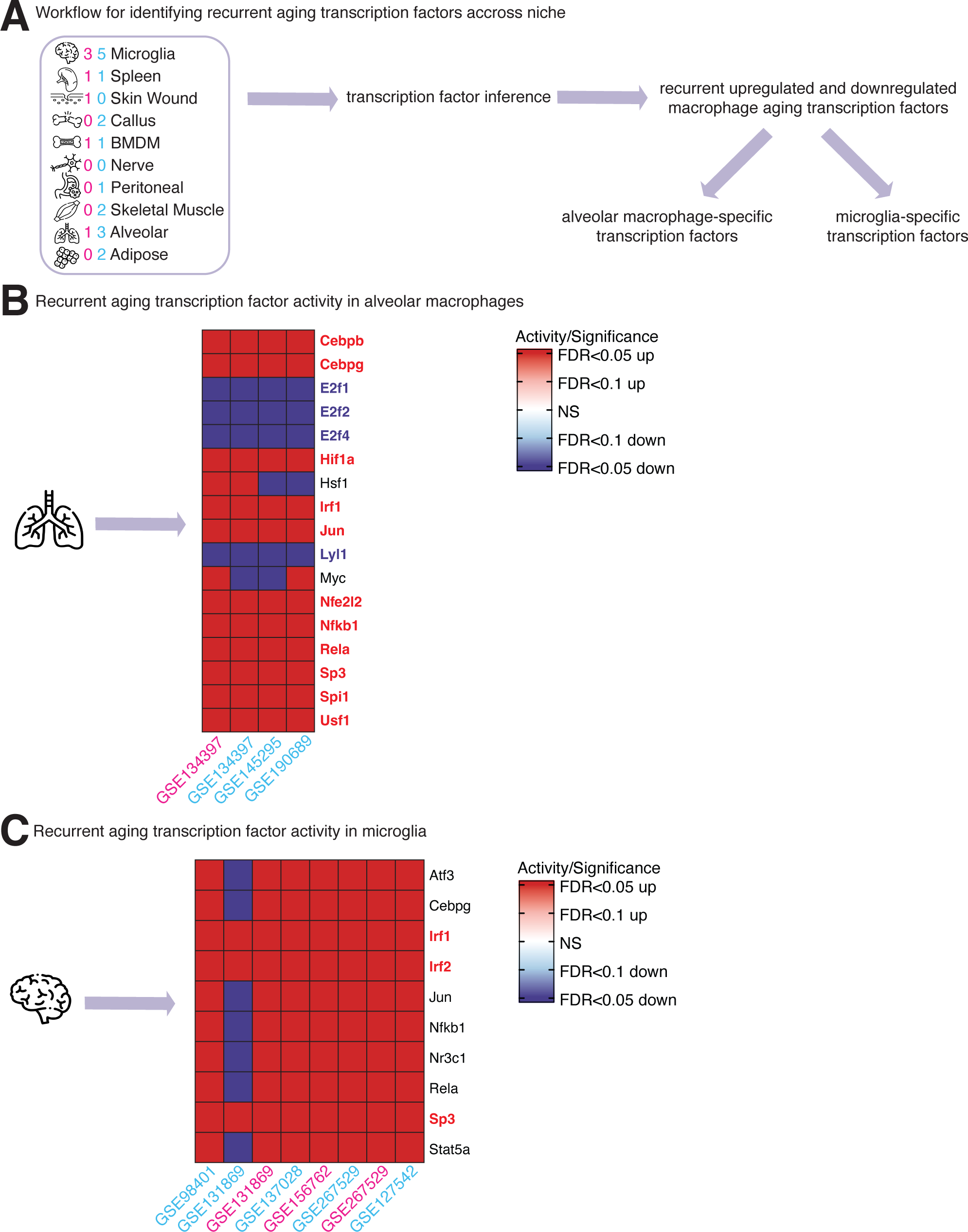
Analysis of predicted changes in transcription factor activity with aging in alveolar macrophages and microglia. **(A)** Schematic overview of transcription factor activity inference analysis for alveolar macrophages and microglia. **(B)** Recurrent transcription factor activity in alveolar macrophage datasets based on decoupleR predictions. **(C)** Recurrent transcription factor activity in microglia datasets based on decoupleR predictions.

Age-related activation of these transcription factor families is well supported in literature, and consistent with known aging phenotypes. For instance, Atf4 activation is strongly linked to neurodegenerative disease via ER stress pathways [104, 105]. Previous chromatin accessibility analyses of aging microglia have demonstrated widespread remodeling of the transcription factor regulatory landscape, with increased accessibility at genomic regions enriched for binding motifs of Irf1, Irf2, Cebpb, and PU.1 with age in mice [106]. The coordinated downregulation of E2f transcription factors and upregulation of stress-responsive Atf, Irf, and Cebp families suggests that tissue-resident macrophages undergo a transcriptional reprogramming characterized by reduced cell-cycle activity and enhanced engagement of stress-related and inflammatory pathways with age.

### Niche-specific TF-pathway networks reveal divergent transcriptional programs underlying alveolar macrophage and microglial aging

Finally, using the same network-level analysis approach as previously described, we examined whether transcription factors with age-altered predicted activity in alveolar macrophages and microglia converge on niche-specific pathways **(Figure 10A)**. In alveolar macrophages, E2F family transcription factors showed the greatest overlap with target genes enriched in downregulated cell cycle-related pathways **(Figure 10B)**, consistent with the predicted age-associated decline in E2F activity observed in this niche and the known role of E2F family members in regulating G1/S transition and DNA synthesis. In microglia, transcription factor targets were most enriched in pathways related to antiviral defense, antibacterial responses, and cytokine production **(Figure 10C)**, consistent with the broadly heightened immune activation signature observed in aging microglia and the predicted increased activity of stress-responsive transcription factors such as Atf3, Irf1, and Irf2 in this niche. Together, these niche-specific network patterns suggest that while alveolar macrophage aging is characterized by decreased proliferation on a transcriptional level, microglial aging may be more influenced by a coordinated engagement of immune effector gene programs.

**Figure 10.**
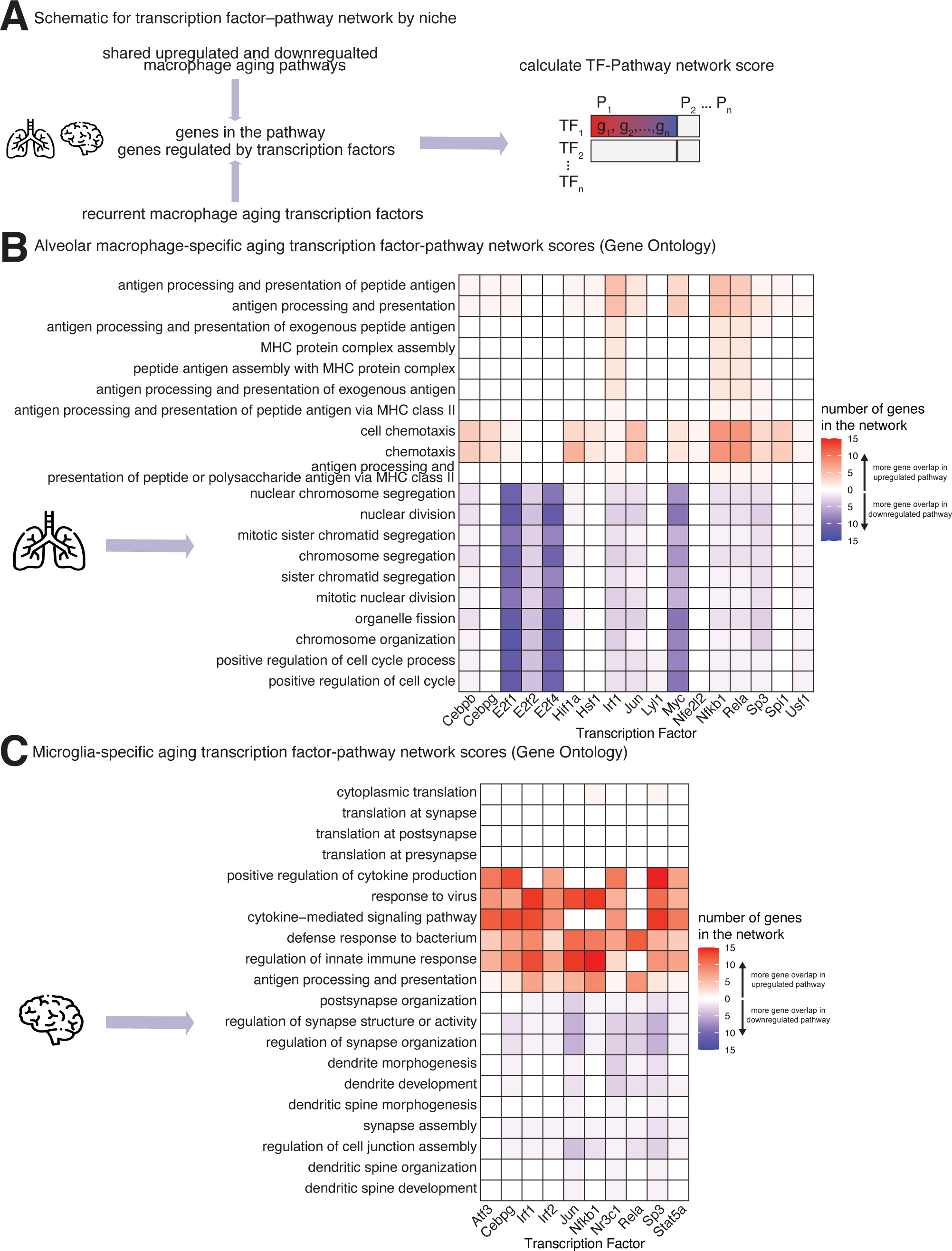
Transcription factor-gene-pathway network for niche-specific transcription factors and niche-specific common upregulated and downregulated macrophage aging signatures. **(A)** Schematic overview of network level analysis between niche-specific enriched transcription factors and niche-specific macrophage aging signatures shared between the sexes. **(B)** Heatmap showing the number of genes in the transcription factor-pathway network for alveolar macrophage-specific Gene Ontology Biological Process (GOBP) pathways upregulated or downregulated with age across both sexes. **(C)** Heatmap showing the number of genes in the transcription factor-pathway network for microglia-specific Gene Ontology Biological Process (GOBP) pathways upregulated or downregulated with age across both sexes.

## DISCUSSION

### Conserved tissue resident macrophage aging signatures

Tissue-resident macrophages are specialized immune cells that reside permanently within various tissues throughout the body. Unlike circulating monocytes that are recruited during inflammation, these macrophages develop from embryonic precursors and are maintained locally under steady-state conditions. They serve critical roles in tissue homeostasis, including clearance of cellular debris, tissue remodeling, and the orchestration of immune responses tailored to their specific microenvironment. Their phenotype and function are finely tuned by the local niche they inhabit, allowing them to adapt to the unique physiological demands and challenges of each tissue [11, 13]. The transcriptomic changes that occur among murine tissue-resident macrophages with age are highly niche-specific. Similar to development and function, the aging of this immune cell type is likely influenced by its microenvironment and exposure to circulating factors throughout the organism’s lifespan [107]. In fact, there is some evidence that peripheral immune organs can affect immunosenescence of immune cells [108]. Overall, we observed higher similarity of differentially expressed genes in among microglia datasets and alveolar macrophage datasets. This could be because of true transcriptomic similarity with aging or increased statistical power. Due to the higher number of datasets for these niches, we were able to do niche-specific analysis for both alveolar macrophages and microglia. On a niche-level, there are notable sex-specific differences in enriched pathways. The upregulation of chemotaxis, antigen presentation, and cytokine production in both sexes and niches aligns with previous reports of age-associated inflammatory signaling in macrophages [12, 109].

In our analysis, we found that the negative regulation of ferroptosis is upregulated across niches. The role of ferroptosis in aging and macrophage biology has been documented, with evidence that ferroptosis is progressively amplified with age [53, 110, 111]. This recently discovered type of cell death is characterized by the overload of iron and membrane lipid peroxidation [112]. Iron is known to accumulate with age in cells [113], especially in the brain [114, 115], and induce pro-inflammatory macrophage polarization [116, 117]. Ferroptosis has been also implicated in the development of autoimmune disease [54]. This upregulation of genes related to *Nfe2l2*-mediated antioxidant signaling across macrophage niches likely represents a compensatory response to age-related iron overload and its associated lipid peroxidation. This may also drive inflammaging indirectly by differentially tuning macrophage polarization states *in vivo*. The decline in Wnt signaling with age across niches is consistent with previous reports. In aging brains, alterations in Wnt signaling have been widely documented [118, 119], and Wnt signaling can directly influence macrophage polarization [59].

Several transcription factors identified in this analysis have roles in immune aging. Both AP-1 and PU.1 are also known to play a role in “inflammaging” in the murine kidney and liver [120]. Karakaslar *et al.* found that aging is accompanied by a conserved activation of the AP-1 transcription factor complex, including *Fos* and *Jun*, across multiple tissues in C57BL/6J mice, as evidenced by changes in gene expression and chromatin accessibility [121]. The role of Egr1 in aging is multifaceted. Overexpression of the transcription factor in *C. elegans* promotes lifespan extension, and decreased expression shortens lifespan. Overall, the expression of Egr1 tends to increase with age and may promote stress adaptive survival [122]. Finally, we note that transcription factor inference algorithms cannot reliably distinguish between closely related family members with overlapping binding motifs [123]. Thus, we recommend caution in interpretations that are part of large closely related families (E2F, CEBP).

Although the role of *Mapk14* in aging remains relatively unexplored, its downregulation in murine macrophages with age may indicate suppression of anti-inflammatory signaling. In one study, reduced CSF-1R expression was observed only in splenic macrophages from aged mice, suggesting cell type and tissue-specific differences in CSF-1R signaling [87]. *Nrros* is downregulated with age in macrophages across niches, meaning that the ability of macrophages to limit the generation of ROS induced by inflammatory signaling may be depreciated with age. Interestingly, serum levels of IL-18BP are elevated in centenarians compared to younger controls [89], and its increased expression alongside the proinflammatory cytokine IL-18 suggests that IL-18BP may serve a protective role against “inflammaging” [124]. The inclusion of two uncharacterized genes (*C130026I21Rik* and *A530032D15Rik*) within this conserved core set further suggests that previously unstudied genes may also contribute to shared macrophage aging processes.

### Alveolar macrophage specific transcriptomic aging signatures

In alveolar macrophages, male-specific enrichment for cytokine production corresponds well with evidence that males experience more pro-inflammatory shifts and heightened innate immune activation with age, including increased cytokine production [51]. The upregulation of genes related to defense response to bacteria in male alveolar macrophages could be related to the fact that male mice tend to have decreased bacteria clearance and survival from bacterial pneumonia [125, 126]. Interestingly, cytoplasmic translation genes were upregulated in male alveolar macrophages and downregulated in female alveolar macrophages with age, suggesting that translation dysregulation may be more sex-dimorphic in the context of alveolar macrophage aging. Downregulation of androgen signaling and sterol-related pathways in males is consistent with studies suggesting sex hormones influence macrophage aging and function [51, 127].

### Microglia-specific transcriptomic aging signatures

Microglia-specific downregulation of genes involved in actin cytoskeleton organization, dendritic spine morphology, and synaptic function aligns with reports implicating microglial actin cytoskeleton regulation in cognitive dysfunction [128, 129]. Interestingly, Rho GTPases, specifically Rho A, are emerging as key players in the regulation of microglial actin dynamics and are implicated in the pathogenesis of neurodegenerative disease [130]. The female-specific aging signature, characterized by the upregulation of signaling pathways related to ER stress and proteostasis, is relatively underreported in the literature in the context of aging. However, one study found that microglia from female mice mount a stronger ER stress response following traumatic brain injury, suggesting that microglial ER stress signaling may differ between sexes in response to injury or aging [131].

## CONCLUSIONS

Collectively, these observations indicate that macrophage aging is not uniform across tissues and that distinct regulatory networks may drive age-associated phenotypes in different tissue-resident macrophage populations. Tissue resident macrophages exhibit diverse regulatory mechanisms shaping immune function across the lifespan. Future aging research should incorporate macrophages from diverse tissue niches, rather than focusing solely on easily accessible or well-studied populations, to better capture the biological heterogeneity underlying macrophage aging.

## LIMITATIONS

It is important to acknowledge several limitations of this meta-analysis, most of which inherent to the reliance of this work on previously generated and published datasets by the scientific community. First, the 24 datasets analyzed here combine publicly available RNA-seq data that encompass non-trivial variations in experimental design, library preparation, sequencing parameters, and sequencing technology due to the original researchers’ aims and experimental designs. We also want to note that there is a wide range in the number of samples across studies (4 to 42). This is known to impact discovery power in differential gene expression analysis and would influence cross study comparisons [132]. Due to these features, it is likely that our discovery power is limited, thus leading to the existing of many false negatives (*i.e*. genes that are actually changed across macrophage niches with age, but that we lack the power to support as such). Importantly, our meta-analysis framework helps reduce the impact of false negatives by leveraging the combined power of many independent studies – and also reduces the odds of including false positive genes that would not get consistent support across many studies.

To note, the datasets we included also differ in (i) the age timepoints sampled (with ‘young’ spanning 2–6 months and ‘old’ spanning 10-26 months), and (ii), in some cases, even the specific macrophage subpopulation profiled within a given tissue (*e.g*., VAT versus eWAT in adipose, or red pulp *vs*. total splenic macrophages), both of which may contribute to dataset-level heterogeneity beyond aging effects

While these datasets provide a valuable foundation for exploring aging-associated transcriptomic changes in macrophages across tissue niches in this meta-analysis study, future standalone studies using data generated under standardized conditions and systematically integrating both sexes would likely provide additional insights without interference from potential batch effects [133]. Indeed, variations in animal (sub)strains, diets, microbiota, library preparation chemistry, sequencing platforms, and inter-laboratory practices introduce substantial sources of technical variability that can sometime mask important but more subtle biological signals, such as those occurring with physiological aging [133]. Additionally, this work focuses primarily on the macrophage aging transcriptome, leaving aging-related changes in the genome, epigenome, and proteome unexplored. Comparative analyses in human macrophage models will further strengthen the translational relevance of these results and advance understanding of macrophage aging in the context of human health and disease.

## METHODS

### Data acquisition

Publicly available datasets were identified through (i) literature review of studies profiling macrophages or microglia from young and aged *Mus musculus* samples, and (ii) targeted searches on GEO and SRA. Datasets were retained if they met the following criteria: originated from *Mus musculus*, used either bulk or single-cell RNA sequencing, profiled niche-specific macrophages or microglia, and included at least two distinct age groups. All datasets included and their corresponding sequencing chemistry are listed in **Supplementary Table S1**. All bulk RNA-seq datasets were downloaded and processed using a unified preprocessing pipeline.

### RNA-seq analysis pipeline

“Read lengths in each dataset range from 50 bp to 150 bp and both single– and paired-end libraries are represented. To harmonize across studies, all reads underwent adapter and quality trimming using Fastx Trimmer. Reads over 75 bp were trimmed to 75 bp prior to alignment. Reads originally shorter than 75 bp were quality and adapter trimmed only to retain their remaining length. Trimmed reads were mapped to the mm10 genome reference using STAR (v.2.7.11b). STAR was run in single– or paired-end mode as appropriate for each library. Read counts were assigned to genes from the UCSC mm10 reference using Subread (v.2.0.2)[134] Gene-level read count tables were imported into R (version 4.5.0) to perform downstream analyses. Only genes with mapped reads in at least one RNA-seq library in each individual dataset were retained for downstream analysis. Count matrices were filtered to remove zero-count genes, then processed with the ‘sva’ package (v3.54.0)[135] to estimate surrogate variables; when present, these were regressed out using ‘removeBatchEffect’ from the ‘limma’ package (v3.62.2)[136]. To ensure reproducibility of surrogate variable estimation and downstream differential expression results, a seed was set in R prior to processing each dataset. DESeq2 (v1.48.2) [137] was used to normalize raw read counts. Genes with FDR <5% were considered significant and are reported in **Supplementary Table S2**.

### Single-cell RNA-seq analysis pipeline

Single-cell RNA sequencing (scRNA-seq) data for two skeletal muscle macrophage datasets were obtained from the NCBI GEO accession GSE205395 and GSE195507. Raw data files in HDF5 format were downloaded and processed through a uniform preprocessing pipeline **(Supplementary Figure 8A)**. Initial quality control was performed to remove low-quality cells. Cells were retained if they expressed more than 250 genes, had fewer than 50,000 total transcripts, and contained less than 25% mitochondrial-derived RNA. Following QC, data were normalized using SCTransform from Seurat (v5.3.0) [138] and glmGamPoi (v1.20.0), with regression of technical covariates including gene count, total RNA count, and mitochondrial content.

To select for macrophages *in silico*, cells expressing canonical macrophage markers *Itgam* (encoding Cd11b) and *Adgre1* (encoding F4/80) were selected as *<u>bona fide</u>* macrophages for downstream analysis. Gene expression profiles from the filtered macrophage population were then aggregated into pseudobulk profiles at the sample level, resulting in one pseudobulk profile per biological replicate and condition. Raw pseudobulk counts were exported for differential gene expression analysis.

### Quality Control pipeline for dataset retention

Only datasets with at least two samples per age group were included in the analysis. Samples from mixed or unknown sex datasets were assigned to male or female groups based on *Xist* and *Ddx3y* expression. Sample ages were extracted from metadata labeling conventions and modeled as a continuous variable for all downstream analyses. Samples with total read depth <1 million reads were removed, and datasets with >50% read depth imbalance between age groups were flagged. Variance-stabilized counts were obtained with DESeq2 (v1.48.2), which was also used for differential expression analysis with age modeled as a continuous variable. Visual QC included MDS plots, gene expression boxplots, and *Xist*/*Ddx3y* h to verify sample sex labels were correctly assigned.

### Jaccard index analysis

For each dataset, the top 1500 upregulated and top 1500 downregulated genes were selected based on positive or negative log_2_FC change values. Pairwise Jaccard indices were then calculated between these gene sets across all datasets. The Jaccard index measures the similarity between two sets and is defined as the size of the intersection divided by the size of the union of the sets. Separate Jaccard similarity matrices were generated for the upregulated and downregulated gene sets to quantify the overlap of differentially expressed genes between datasets. The jaccard index similarity scores were used to create a heatmap showing similarity between each dataset.

### Overrepresentation analysis

Over-representation analysis (ORA) was conducted using the R package clusterProfiler (v4.16.0) [139] along with the Bioconductor annotation package org.Mm.eg.db (v3.21.0) for Gene Ontology annotations. For each macrophage niche, differentially expressed genes (DEGs) shared across datasets were separated into upregulated and downregulated groups based on the sign of the DESeq2 log_2_(fold change). These gene lists served as the input for enrichment, while the background (universe) was defined as all genes expressed across datasets within the same niche. Gene Ontology (GO) enrichment was performed using the enrichGO function, restricting results to biological process (BP) terms.

Visualizations were created using the ggplot2 package (v4.0.0) [140], and all significant enrichment results are provided in **Supplementary Table 7.**

### Multivariate gene set enrichment analysis

To identify biological pathways commonly regulated across macrophage populations from different tissues and conditions, we used the R package mitch (v1.20.0) [52], which allows for the simultaneous analysis of multiple gene expression datasets in a single gene set enrichment analysis. Differentially expressed gene sets were mapped to Reactome [141], KEGG and Gene Ontology Biological Process (GO:BP) pathway gene sets curated in the Molecular Signatures Database (MSigDB), accessed via the msigdbr R package (v.25.1.1) [142]. To determine whether pathways were consistently upregulated or downregulated across datasets, we calculated a consensus direction of enrichment for each pathway using the signed enrichment scores. Terms were defined as consistently upregulated or downregulated if they showed the same sign of enrichment in at least 18/24 (75%) macrophage datasets. Enrichment was prioritized by significance and then terms were ranked by absolute value of enrichment score for a given pathway across datasets to capture significantly enriched pathways with the greatest effect size.

### Meta-analysis to identify core macrophage aging genes

To identify core macrophage aging genes consistently differentially expressed with age across multiple transcriptomic datasets, we conducted a meta-analysis using the metaRNASeq R package (v1.0.8) [81]. Genes flagged by DESeq2’s independent filtering, indicated by an NA adjusted p-value, were removed prior to meta-analysis. This step ensured that the distribution of raw p-values was approximately uniform for each dataset as required by the p-value combination methods implemented in metaRNASeq. We identified a common gene universe of 9,225 genes and each dataset was then reduced to include only these common genes. To assess the consistency of differential expression direction, we calculated the sign of the log_2_FC (positive or negative) for each gene across studies and computed the number of datasets with a positive log_2_FC and negative log_2_FC for that gene. Genes were considered to have consistent directionality if at least 18 out of 24 datasets showed the same sign. Genes were defined as consistently differentially expressed if they were detected as significantly differentially expressed in either Fisher or inverse normal meta-analysis (adjusted p ≤ 0.05) and showed consistent directionality of log_2_FC in at least 18/24 datasets. Adjusted meta-analysis p-values were computed using the Benjamini–Hochberg (BH) method. These genes were retained as the high-confidence meta-analysis derived macrophage aging genes.

Over-representation analysis (ORA) was conducted for Gene Ontology (GO), Reactome, and KEGG pathway databases using the clusterProfiler and ReactomePA R packages. GO enrichment was performed using the enrichGO function with the ontology parameter set to “ALL” to include Biological Process (BP), Molecular Function (MF), and Cellular Component (CC) categories in a single analysis. Reactome pathway enrichment was performed using the enrichPathway function from the ReactomePA package, and KEGG pathway enrichment was performed using the enrichKEGG function from clusterProfiler. For all three databases, enrichment was tested separately for consistently upregulated and consistently downregulated genes, using a custom background gene set defined as the universe of expressed genes common across all macrophage datasets. Gene identifiers were mapped to Entrez IDs using the Mus musculus annotation database (org.Mm.eg.db). P-values were adjusted for multiple testing using the Benjamini–Hochberg method, and terms with adjusted p-values below 0.05 were considered statistically significant.

### Transcription factor activity inference analysis

To infer transcription factor (TF) activity changes associated with aging across datasets, we used the decoupleR R package (v2.14.0), which enables the estimation of regulator activity from gene expression data using prior knowledge. Wald statistics from aging comparisons were used as input. We employed the normalized enrichment score (norm_fgsea) method implemented in decoupleR, which uses a gene set enrichment framework to assess the enrichment of TF target gene sets. Prior knowledge of mouse TF-target relationships was obtained via the get_collectri() function, which provides curated transcriptional regulatory networks from the CollecTRI resource [143]. For each dataset, TF activities were inferred separately, and only regulators with significant enrichment at FDR < 5% and < 10% were retained. Recurrently enriched TFs across multiple datasets were identified by filtering those detected in more than half of the datasets. A consensus heatmap was generated to summarize consistent activation or repression of these regulators with age across each macrophage dataset by significance cutoff.

### Transcription factor-gene-pathway network analysis

To integrate transcription factor regulatory relationships with age-associated pathway enrichment, we constructed a TF-gene-pathway network linking inferred TF targets to consistently enriched biological pathways. TF-target gene relationships were obtained from the CollecTRI mouse transcription factor-target regulatory network [143]. Differentially expressed genes were defined as those with an FDR below 0.15; a more permissive threshold was applied to maximize detection power of genes potentially driving Mitch pathway enrichment results, as Mitch evaluates gene-level statistics jointly across multiple contrasts rather than applying a gene-level significance filter directly, ensuring these genes were represented in the downstream TF-gene-pathway network. For each pathway database (Gene Ontology Biological Process, KEGG, and Reactome), gene sets for *Mus musculus* were retrieved using the msigdbr R package (v.25.1.1). Consistently enriched pathways were identified from Mitch multi-dataset enrichment results by requiring that the enrichment score sign was concordant in at least 18 out of 24 datasets (i.e. ≥75% of datasets). The top 10 consistently upregulated and top 10 consistently downregulated pathways were selected for network construction. For each selected pathway, the overlap between pathway member genes and TF target genes was computed. Results were visualized as heatmaps, showing the number of genes (TF targets) enriched in upregulated pathways and downregulated pathways. This analysis was performed using the overall recurrent macrophage aging TF list as well as niche-specific recurrent aging TF lists for alveolar macrophages and microglia, with tissue-specific pathway inputs derived from over-representation analysis (ORA) of sex-shared age-associated genes.

## ACKNOWLEDGEMENTS

This work was supported by Hevolution grant HF-GRO-23-1199072-28 to B.A.B. and USC Leonard Davis School of Gerontology Biology of Aging PhD Research Fellowships to ET.

## AUTHOR CONTRIBUTIONS

**Ella Schwab:** Formal analysis; data curation; writing – original draft; writing – review and editing. **Eyael Tewelde:** Formal analysis; writing – review and editing. **Leon Chen**: Formal analysis; writing – review and editing. **Bérénice A Benayoun:** Conceptualization; data curation; supervision; funding acquisition; writing – review and editing.

## CODE AVAILABILITY

All analytical scripts for this study are available on the Benayoun Lab Github https://github.com/BenayounLaboratory/ATOM.

## Legends to Supplementary Figures

**Supplementary Figure S1:**

Sex marker gene expression across datasets stratified by author-reported and QC-annotated sex labels. **(A)** Heatmaps showing VST-normalized expression of *Xist* and *Ddx3y* across datasets, shown alongside author-reported and QC-annotated sex. Red stars indicate samples or datasets that had unclear sex and did not pass this QC step.

**Supplementary Figure S2**:

Jaccard index representing transcriptomic similarity score between datasets for genes differentially expressed with age (FDR < 5%). **(A)** Similarity scores of genes significantly upregulated with age and **(B)** significantly downregulated with age across niche.

**Supplementary Figure S3**:

Multivariate gene set enrichment analysis of age-associated pathway enrichment across macrophage niches using the KEGG database. Pathways upregulated with age **(A)** and downregulated with age **(B)** are shown.

**Supplementary Figure S4**:

Additional transcription factor-pathway networks. **(A)** Heatmap showing the number of genes in the transcription factor-pathway network for upregulated and downregulated KEGG pathways with age across macrophage niches.

**Supplementary Figure S5**:

Sensitivity analysis for different consistency thresholds and differentially expressed genes in 21 out of 24 datasets. **(A)** Bar plot showing the sensitivity analysis for p-value combination derived DEGs based on different consistency thresholds. **(B)** Gene expression heatmap displaying a core set of age-associated macrophage genes consistent in at least 21/24 datasets across all macrophage niches, colored by log_2_ fold change (log_₂_FC) with age. Positive log_2_ fold change (log_2_FC) values indicate increased expression with age, and negative log_2_FC values indicate decreased expression with age.

**Supplementary Figure S6**:

Upset plots showing conserved male and female murine macrophage aging genes within each niche (FDR < 5%). Intersection set sizes for differentially expressed aging genes in **(A)** spleen macrophages, **(B)** callus macrophages, **(C)** skeletal muscle macrophages, **(D)** bone-marrow derived macrophages, and **(E)** adipose macrophages

**Supplementary Figure S7**:

Transcription factor pathway gene network for niche-specific transcription factors and common upregulated and downregulated macrophage aging pathway-level signatures. **(A)** Schematic overview of network level analysis between niche-specific enriched transcription factors and pathway level macrophage aging signatures. **(B)** Heatmap showing the number of genes in the transcription factor-pathway network for alveolar macrophage-specific Gene Ontology Biological Process (GOBP), Reactome, and KEGG pathways upregulated or downregulated with age across all niches. **(C)** Heatmap showing the number of genes in the transcription factor-pathway network for microglia-specific Gene Ontology Biological Process (GOBP), Reactome, and KEGG pathways upregulated or downregulated with age across all niches.

**Supplementary Figure S8**:

Single-cell analysis workflow. **(A)** Workflow diagram describing analysis pipeline for two single cell datasets GSE195507 and GSE205395 including quality control, *in silico* purification, and pseudobulking.

## Inventory of Supplementary Tables

**Supplementary Table S1:**

(A) Table depicting quality control workflow and associated metadata for each macrophage aging dataset. (B) RNA extraction and enrichment method, sequencing chemistry, type, platform, read length, strandedness, and mouse strains for each macrophage aging dataset.

**Supplementary Table S2**:

1-29: DESeq2 output for each dataset

**Supplementary Table S3**:

(A) Top 1500 genes downregulated with age across all niches. (B) Top 1500 genes upregulated with age across all niches. (C) List of all genes downregualted with age (FDR < 5%). (D) List of all genes upregualted with age (FDR < 5%).

**Supplementary Table S4**:

(A) Gene Ontology biological process aging pathways ranked by effect size. (B) Gene Ontology biological process aging pathways ranked by significance. (C) Reactome aging pathways ranked by effect size. (D) Reactome aging pathways ranked by significance. (E) KEGG aging pathways ranked by effect size. (D) KEGG aging pathways ranked by significance.

**Supplementary Table S5**:

(A) List of predicted transcription factor activity changing with age detected by DecoupleR.

**Supplementary Table S6**:

(A) List of 593 Conserved murine macrophage aging genes consistent in at least 18/24 datasets. (B) List of 35 Conserved murine macrophage aging genes consistent in at least 21/24 datasets. (C) Number of meta-analysis DEGs per threshold from sensitivity analysis. (D) ORA results for 342 upregulated genes from the Gene Ontology database. (E) ORA results for 342 upregulated genes from the Reactome database. (F) ORA results for 342 upregulated genes from the KEGG database. (G) ORA results for 251 downregulated genes from the Gene Ontology database. (H) ORA results for 251 downregulated genes from the Reactome database. (I) ORA results for 251 downregulated genes from the KEGG database.

**Supplementary Table S7**:

(A) List of genes upregulated in alveolar macrophages with age (FDR < 5%). (B) List of genes downregulated in alveolar macrophages with age (FDR <5%). (C) List of genes upregulated in microglia with age (FDR < 5%). (D) List of genes downregulated in microglia with age (FDR <5%)

## REFERENCES

1. Peters MJ, Joehanes R, Pilling LC, Schurmann C, Conneely KN, Powell J, Reinmaa E, Sutphin GL, Zhernakova A, Schramm K et al: The transcriptional landscape of age in human peripheral blood. Nature Communications 2015, 6(1):8570.

2. Palmer D, Fabris F, Doherty A, Freitas AA, de Magalhães JP: Ageing transcriptome meta-analysis reveals similarities and differences between key mammalian tissues. Aging 2021, 13(3):3313–3341.

3. López-Otín C, Blasco MA, Partridge L, Serrano M, Kroemer G: Hallmarks of aging: An expanding universe. Cell 2023, 186(2):243–278.

4. Schaum N, Lehallier B, Hahn O, Pálovics R, Hosseinzadeh S, Lee SE, Sit R, Lee DP, Losada PM, Zardeneta ME et al: Ageing hallmarks exhibit organ-specific temporal signatures. Nature 2020, 583(7817):596–602.

5. Glass D, Viñuela A, Davies MN, Ramasamy A, Parts L, Knowles D, Brown AA, Hedman ÅK, Small KS, Buil A et al: Gene expression changes with age in skin, adipose tissue, blood and brain. Genome Biology 2013, 14(7):R75.

6. Yamamoto R, Chung R, Vazquez JM, Sheng H, Steinberg PL, Ioannidis NM, Sudmant PH: Tissue-specific impacts of aging and genetics on gene expression patterns in humans. Nature Communications 2022, 13(1):5803.

7. Zhang MJ, Pisco AO, Darmanis S, Zou J: Mouse aging cell atlas analysis reveals global and cell type-specific aging signatures. ELife 2021, 10.

8. Almanzar N, Antony J, Baghel AS, Bakerman I, Bansal I, Barres BA, Beachy PA, Berdnik D, Bilen B, Brownfield D et al: A single-cell transcriptomic atlas characterizes ageing tissues in the mouse. Nature 2020, 583(7817):590–595.

9. Hammond TR, Dufort C, Dissing-Olesen L, Giera S, Young A, Wysoker A, Walker AJ, Gergits F, Segel M, Nemesh J et al: Single-Cell RNA Sequencing of Microglia throughout the Mouse Lifespan and in the Injured Brain Reveals Complex Cell-State Changes. Immunity 2019, 50(1):253–271.e256.

10. Rando TA, Wyss-Coray T: Asynchronous, contagious and digital aging. Nature Aging 2021, 1(1):29–35.

11. Guan F, Wang R, Yi Z, Luo P, Liu W, Xie Y, Liu Z, Xia Z, Zhang H, Cheng Q: Tissue macrophages: origin, heterogenity, biological functions, diseases and therapeutic targets. Signal Transduction and Targeted Therapy 2025, 10(1):93.

12. Wynn TA, Chawla A, Pollard JW: Macrophage biology in development, homeostasis and disease. Nature 2013, 496(7446):445–455.

13. Wu Y, Hirschi KK: Tissue-Resident Macrophage Development and Function. Frontiers in Cell and Developmental Biology 2021, 8.

14. Gomez Perdiguero E, Klapproth K, Schulz C, Busch K, Azzoni E, Crozet L, Garner H, Trouillet C, de Bruijn MF, Geissmann F et al: Tissue-resident macrophages originate from yolk-sac-derived erythro-myeloid progenitors. Nature 2015, 518(7540):547–551.

15. Lazarov T, Juarez-Carreño S, Cox N, Geissmann F: Physiology and diseases of tissue-resident macrophages. Nature 2023, 618(7966):698–707.

16. Gautier EL, Shay T, Miller J, Greter M, Jakubzick C, Ivanov S, Helft J, Chow A, Elpek KG, Gordonov S et al: Gene-expression profiles and transcriptional regulatory pathways that underlie the identity and diversity of mouse tissue macrophages. Nature Immunology 2012, 13(11):1118–1128.

17. Lavin Y, Winter D, Blecher-Gonen R, David E, Keren-Shaul H, Merad M, Jung S, Amit I: Tissue-Resident Macrophage Enhancer Landscapes Are Shaped by the Local Microenvironment. Cell 2014, 159(6):1312–1326.

18. Liu Z, Liang Q, Ren Y, Guo C, Ge X, Wang L, Cheng Q, Luo P, Zhang Y, Han X: Immunosenescence: molecular mechanisms and diseases. Signal Transduction and Targeted Therapy 2023, 8(1):200.

19. Jackaman C, Tomay F, Duong L, Abdol Razak NB, Pixley FJ, Metharom P, Nelson DJ: Aging and cancer: The role of macrophages and neutrophils. Ageing research reviews 2017, 36:105–116.

20. Inomata M, Xu S, Chandra P, Meydani SN, Takemura G, Philips JA, Leong JM: Macrophage LC3-associated phagocytosis is an immune defense against *Streptococcus pneumoniae* that diminishes with host aging. Proceedings of the National Academy of Sciences 2020, 117(52):33561–33569.

21. Thevaranjan N, Puchta A, Schulz C, Naidoo A, Szamosi JC, Verschoor CP, Loukov D, Schenck LP, Jury J, Foley KP et al: Age-Associated Microbial Dysbiosis Promotes Intestinal Permeability, Systemic Inflammation, and Macrophage Dysfunction. Cell Host & Microbe 2017, 21(4):455–466.e454.

22. Dube CT, Ong YHB, Wemyss K, Krishnan S, Tan TJ, Janela B, Grainger JR, Ronshaugen M, Mace KA, Lim CY: Age-Related Alterations in Macrophage Distribution and Function Are Associated With Delayed Cutaneous Wound Healing. Frontiers in Immunology 2022, 13.

23. Sebastián C, Espia M, Serra M, Celada A, Lloberas J: MacrophAging: A cellular and molecular review. Immunobiology 2005, 210(2-4):121–126.

24. Oishi Y, Manabe I: Macrophages in age-related chronic inflammatory diseases. npj Aging and Mechanisms of Disease 2016, 2(1):16018.

25. Moss CE, Phipps H, Wilson HL, Kiss-Toth E: Markers of the ageing macrophage: a systematic review and meta-analysis. Frontiers in Immunology 2023, 14.

26. Hao D, Caja KR, McBride MA, Owen AM, Bohannon JK, Hernandez A, Ali S, Dalal S, Williams DL, Sherwood ER: Trained immunity enhances host resistance to infection in aged mice. Journal of leukocyte biology 2025, 117(4).

27. McGill CJ, White OS, Lu RJ, Sampathkumar NK, Benayoun BA: Sex-dimorphic gene regulation in murine macrophages across niches. Immunology & Cell Biology 2025, 103(6):563–577.

28. Seyfried AN, McCabe A, Smith JNP, Calvi LM, MacNamara KC: CCR5 maintains macrophages in the bone marrow and drives hematopoietic failure in a mouse model of severe aplastic anemia. Leukemia: official journal of the Leukemia Society of America, Leukemia Research Fund, UK 2021, 35(11):3139–3151.

29. Wang Y, Deng W, Lee D, Yan L, Lu Y, Dong S, Huntoon K, Antony A, Li X, Ye R et al: Age-associated disparity in phagocytic clearance affects the efficacy of cancer nanotherapeutics. Nature nanotechnology 2024, 19(2):255–263.

30. Deczkowska A, Matcovitch-Natan O, Tsitsou-Kampeli A, Ben-Hamo S, Dvir-Szternfeld R, Spinrad A, Singer O, David E, Winter DR, Smith LK et al: Mef2C restrains microglial inflammatory response and is lost in brain ageing in an IFN-I-dependent manner. Nature Communications 2017, 8(1).

31. Gyoneva S, Hosur R, Gosselin D, Zhang B, Ouyang Z, Cotleur AC, Peterson M, Allaire N, Challa R, Cullen P et al: Cx3cr1-deficient microglia exhibit a premature aging transcriptome. Life Science Alliance 2019, 2(6):e201900453.

32. Kang S, Ko EY, Andrews AE, Shin JE, Nance KJ, Barman PK, Heeger PS, Freeman WM, Benayoun BA, Goodridge HS: Microglia undergo sex-dimorphic transcriptional and metabolic rewiring during aging. Journal of Neuroinflammation 2024, 21(1).

33. Keane L, Antignano I, Riechers S-P, Zollinger R, Dumas AA, Offermann N, Bernis ME, Russ J, Graelmann F, McCormick PN et al: mTOR-dependent translation amplifies microglia priming in aging mice. Journal of Clinical Investigation 2021, 131(1).

34. Pan J, Ma N, Yu B, Zhang W, Wan J: Transcriptomic profiling of microglia and astrocytes throughout aging. Journal of Neuroinflammation 2020, 17(1).

35. Pluvinage JV, Haney MS, Smith BAH, Sun J, Iram T, Bonanno L, Li L, Lee DP, Morgens DW, Yang AC et al: CD22 blockade restores homeostatic microglial phagocytosis in ageing brains. Nature 2019, 568(7751):187–192.

36. Angelidis I, Simon LM, Fernandez IE, Strunz M, Mayr CH, Greiffo FR, Tsitsiridis G, Ansari M, Graf E, Strom T-M et al: An atlas of the aging lung mapped by single cell transcriptomics and deep tissue proteomics. Nature Communications 2019, 10(1).

37. McElroy GS, Chakrabarty RP, D’Alessandro KB, Hu Y-S, Vasan K, Tan J, Stoolman JS, Weinberg SE, Steinert EM, Reyfman PA et al: Reduced expression of mitochondrial complex I subunit Ndufs2 does not impact healthspan in mice. Scientific Reports 2022, 12(1).

38. McQuattie-Pimentel AC, Ren Z, Joshi N, Watanabe S, Stoeger T, Chi M, Lu Z, Sichizya L, Aillon RP, Chen C-I et al: The lung microenvironment shapes a dysfunctional response of alveolar macrophages in aging. Journal of Clinical Investigation 2021, 131(4).

39. Watanabe S, Markov NS, Lu Z, Piseaux Aillon R, Soberanes S, Runyan CE, Ren Z, Grant RA, Maciel M, Abdala-Valencia H et al: Resetting proteostasis with ISRIB promotes epithelial differentiation to attenuate pulmonary fibrosis. Proceedings of the National Academy of Sciences 2021, 118(20):e2101100118.

40. Krasniewski LK, Chakraborty P, Cui C-Y, Mazan-Mamczarz K, Dunn C, Piao Y, Fan J, Shi C, Wallace T, Nguyen C et al: Single-cell analysis of skeletal muscle macrophages reveals age-associated functional subpopulations. ELife 2022, 11.

41. Runyan CE, Welch LC, Lecuona E, Shigemura M, Amarelle L, Abdala-Valencia H, Joshi N, Lu Z, Nam K, Markov NS et al: Impaired phagocytic function in CX3CR1+ tissue-resident skeletal muscle macrophages prevents muscle recovery after influenza A virus-induced pneumonia in old mice. Aging Cell 2020, 19(9).

42. Camell CD, Sander J, Spadaro O, Lee A, Nguyen KY, Wing A, Goldberg EL, Youm Y-H, Brown CW, Elsworth J et al: Inflammasome-driven catecholamine catabolism in macrophages blunts lipolysis during ageing. Nature 2017, 550(7674):119–123.

43. Hall BM, Gleiberman AS, Strom E, Krasnov PA, Frescas D, Vujcic S, Leontieva OV, Antoch MP, Kogan V, Koman IE et al: Immune checkpoint protein VSIG4 as a biomarker of aging in murine adipose tissue. Aging Cell 2020, 19(10).

44. Blacher E, Tsai C, Litichevskiy L, Shipony Z, Iweka CA, Schneider KM, Chuluun B, Heller HC, Menon V, Thaiss CA et al: Aging disrupts circadian gene regulation and function in macrophages. Nature Immunology 2022, 23(2):229–236.

45. Gal-Oz ST, Maier B, Yoshida H, Seddu K, Elbaz N, Czysz C, Zuk O, Stranger BE, Ner-Gaon H, Shay T: ImmGen report: sexual dimorphism in the immune system transcriptome. Nature Communications 2019, 10(1).

46. Slusarczyk P, Mandal PK, Zurawska G, Niklewicz M, Chouhan K, Mahadeva R, Jończy A, Macias M, Szybinska A, Cybulska-Lubak M et al: Impaired iron recycling from erythrocytes is an early hallmark of aging. ELife 2023, 12.

47. Clark D, Brazina S, Miclau T, Park S, Hsieh CL, Nakamura M, Marcucio R: Age-related changes to macrophage subpopulations and TREM2 dysregulation characterize attenuated fracture healing in old mice. Aging Cell 2024, 23(9).

48. Clark D, Brazina S, Yang F, Hu D, Hsieh CL, Niemi EC, Miclau T, Nakamura MC, Marcucio R: Age-related changes to macrophages are detrimental to fracture healing in mice. Aging Cell 2020, 19(3).

49. Stratton JA, Holmes A, Rosin NL, Sinha S, Vohra M, Burma NE, Trang T, Midha R, Biernaskie J: Macrophages Regulate Schwann Cell Maturation after Nerve Injury. Cell Reports 2018, 24(10):2561–2572.e2566.

50. Shen X, Wang C, Zhou X, Zhou W, Hornburg D, Wu S, Snyder MP: Nonlinear dynamics of multi-omics profiles during human aging. Nature Aging 2024, 4(11):1619–1634.

51. Márquez EJ, Chung C-h, Marches R, Rossi RJ, Nehar-Belaid D, Eroglu A, Mellert DJ, Kuchel GA, Banchereau J, Ucar D: Sexual-dimorphism in human immune system aging. Nature Communications 2020, 11(1):751.

52. Kaspi A, Ziemann M: mitch: multi-contrast pathway enrichment for multi-omics and single-cell profiling data. BMC Genomics 2020, 21(1):447.

53. Mazhar M, Din AU, Ali H, Yang G, Ren W, Wang L, Fan X, Yang S: Implication of ferroptosis in aging. Cell Death Discovery 2021, 7(1).

54. Lai B, Wu C-H, Wu C-Y, Luo S-F, Lai J-H: Ferroptosis and autoimmune diseases. Frontiers in immunology 2022, 13:916664.

55. Bain CC, Macdonald AS: The impact of the lung environment on macrophage development, activation and function: diversity in the face of adversity. Mucosal Immunology 2022, 15(2):223–234.

56. Van Helden SFG, Anthony EC, Dee R, Hordijk PL: Rho GTPase Expression in Human Myeloid Cells. PLoS ONE 2012, 7(8):e42563.

57. Selman M, Pardo A: Fibroageing: An ageing pathological feature driven by dysregulated extracellular matrix-cell mechanobiology. Ageing research reviews 2021, 70:101393.

58. Marino GE, Weeraratna AT: A glitch in the matrix: Age-dependent changes in the extracellular matrix facilitate common sites of metastasis. Aging and Cancer 2020, 1(1-4):19–29.

59. Abaricia JO, Shah AH, Chaubal M, Hotchkiss KM, Olivares-Navarrete R: Wnt signaling modulates macrophage polarization and is regulated by biomaterial surface properties. Biomaterials 2020, 243:119920.

60. Becker L, Nguyen L, Gill J, Kulkarni S, Pasricha PJ, Habtezion A: Age-dependent shift in macrophage polarisation causes inflammation-mediated degeneration of enteric nervous system. Gut 2017, 67(5):827–836.

61. Dimitrijević M, Stanojević S, Blagojević V, Ćuruvija I, Vujnović I, Petrović R, Arsenović-Ranin N, Vujić V, Leposavić G: Aging affects the responsiveness of rat peritoneal macrophages to GM-CSF and IL-4. Biogerontology 2016, 17(2):359–371.

62. Badia-I-Mompel P, Vélez Santiago J, Braunger J, Geiss C, Dimitrov D, Müller-Dott S, Taus P, Dugourd A, Holland CH, Ramirez Flores RO et al: decoupleR: ensemble of computational methods to infer biological activities from omics data. Bioinformatics Advances 2022, 2(1).

63. Mechta-Grigoriou F, Gerald D, Yaniv M: The mammalian Jun proteins: redundancy and specificity. Oncogene 2001, 20(19):2378–2389.

64. Wisdom R, Johnson RS, Moore C: c-Jun regulates cell cycle progression and apoptosis by distinct mechanisms. The EMBO Journal 1999, 18(1):188–197-197.

65. Stepniak E, Ricci R, Eferl R, Sumara G, Sumara I, Rath M, Hui L, Wagner EF: c-Jun/AP-1 controls liver regeneration by repressing p53/p21 and p38 MAPK activity. Genes & Development 2006, 20(16):2306–2314.

66. Johnson RS, Van Lingen B, Papaioannou VE, Spiegelman BM: A null mutation at the c-jun locus causes embryonic lethality and retarded cell growth in culture. Genes & development 1993, 7(7b):1309–1317.

67. Huber R, Pietsch D, Panterodt T, Brand K: Regulation of C/EBPβ and resulting functions in cells of the monocytic lineage. Cellular signalling 2012, 24(6):1287–1296.

68. Ren Q, Liu Z, Wu L, Yin G, Xie X, Kong W, Zhou J, Liu S: C/EBPβ: The structure, regulation, and its roles in inflammation-related diseases. Biomedicine & pharmacotherapy 2023, 169:115938.

69. Ruffell D, Mourkioti F, Gambardella A, Kirstetter P, Lopez RG, Rosenthal N, Nerlov C: A CREB-C/EBPβ cascade induces M2 macrophage-specific gene expression and promotes muscle injury repair. Proceedings of the National Academy of Sciences 2009, 106(41):17475–17480.

70. Li Y, Chen H, Wang J, Wang J, Niu X, Wang C, Qin D, Li F, Wang Y, Xiong J et al: Inflammation-activated C/EBPβ mediates high-fat diet-induced depression-like behaviors in mice. Frontiers in molecular neuroscience 2022, 15.

71. Heinz S, Benner C, Spann N, Bertolino E, Lin YC, Laslo P, Cheng JX, Murre C, Singh H, Glass CK: Simple Combinations of Lineage-Determining Transcription Factors Prime cis-Regulatory Elements Required for Macrophage and B Cell Identities. Molecular Cell 2010, 38(4):576–589.

72. Qi S, Zhang Y, Kong L, Bi D, Kong H, Zhang S, Zhao C: SPI1-mediated macrophage polarization aggravates age-related macular degeneration. Frontiers in Immunology 2024, 15.

73. Kierdorf K, Prinz M, Geissmann F, Gomez Perdiguero E: Development and function of tissue resident macrophages in mice. Seminars in Immunology 2015, 27(6):369–378.

74. Wu Y, Guo W, Kuang H, Wu X, Trinh TH, Wang Y, Zhao S, Wen Z, Yu T: Pu.1/Spi1 dosage controls the turnover and maintenance of microglia in zebrafish and mammals. In.: eLife Sciences Publications, Ltd; 2025.

75. Jones RE, Andrews R, Holmans P, Hill M, Taylor PR: Modest changes in Spi1 dosage reveal the potential for altered microglial function as seen in Alzheimer’s disease. Scientific Reports 2021, 11(1):14935.

76. Trizzino M, Zucco A, Deliard S, Wang F, Barbieri E, Veglia F, Gabrilovich D, Gardini A: EGR1 is a gatekeeper of inflammatory enhancers in human macrophages. Science Advances 2021, 7(3):eaaz8836.

77. Peyssonnaux C, Datta V, Cramer T, Doedens A, Theodorakis EA, Gallo RL, Hurtado-Ziola N, Nizet V, Johnson RS: HIF-1α expression regulates the bactericidal capacity of phagocytes. Journal of Clinical Investigation 2005, 115(7):1806–1815.

78. Cramer T, Yamanishi Y, Clausen BE, Förster I, Pawlinski R, Mackman N, Haase VH, Jaenisch R, Corr M, Nizet V et al: HIF-1α Is Essential for Myeloid Cell-Mediated Inflammation. Cell 2003, 112(5):645–657.

79. Qiu B, Yuan P, Du X, Jin H, Du J, Huang Y: Hypoxia inducible factor-1α is an important regulator of macrophage biology. Heliyon 2023, 9(6):e17167.

80. Luan B, Yoon Y-S, Le Lay J, Kaestner KH, Hedrick S, Montminy M: CREB pathway links PGE2 signaling with macrophage polarization. Proceedings of the National Academy of Sciences 2015, 112(51):15642–15647.

81. Rau A, Marot G, Jaffrézic F: Differential meta-analysis of RNA-seq data from multiple studies. BMC Bioinformatics 2014, 15(1):91.

82. Li Z, Jiao Y, Fan EK, Scott MJ, Li Y, Li S, Billiar TR, Wilson MA, Shi X, Fan J: Aging-Impaired Filamentous Actin Polymerization Signaling Reduces Alveolar Macrophage Phagocytosis of Bacteria. The journal of immunology: official journal of the American Association of Immunologists 2017, 199(9):3176–3186.

83. Gourlay CW, Carpp LN, Timpson P, Winder SJ, Ayscough KR: A role for the actin cytoskeleton in cell death and aging in yeast. The journal of cell biology 2004, 164(6):803–809.

84. Murray RZ, Stow JL: Cytokine Secretion in Macrophages: SNAREs, Rabs, and Membrane Trafficking. Frontiers in Immunology 2014, 5.

85. Canovas B, Nebreda AR: Diversity and versatility of p38 kinase signalling in health and disease. Nature Reviews Molecular Cell Biology 2021, 22(5):346–366.

86. Lira-Junior R, Åkerman S, Gustafsson A, Klinge B, Boström EA: Colony stimulating factor-1 in saliva in relation to age, smoking, and oral and systemic diseases. Scientific Reports 2017, 7(1).

87. Duong L, Pixley FJ, Nelson DJ, Jackaman C: Aging Leads to Increased Monocytes and Macrophages With Altered CSF-1 Receptor Expression and Earlier Tumor-Associated Macrophage Expansion in Murine Mesothelioma. 2022(2673-6217 (Electronic)).

88. Barth E, Srivastava A, Stojiljkovic M, Frahm C, Axer H, Witte OW, Marz M: Conserved aging-related signatures of senescence and inflammation in different tissues and species. Aging 2019, 11(19):8556–8572.

89. Gangemi S, Basile G, Merendino RA, Minciullo PL, Novick D, Rubinstein M, Dinarello CA, Balbo CL, Franceschi C, Basili S et al: Increased circulating Interleukin-18 levels in centenarians with no signs of vascular disease: another paradox of longevity? Experimental Gerontology 2003, 38(6):669–672.

90. Lee JC, Laydon JT, McDonnell PC, Gallagher TF, Kumar S, Green D, McNulty D, Blumenthal MJ, Keys JR, Land vatter SW et al: A protein kinase involved in the regulation of inflammatory cytokine biosynthesis. Nature 1994, 372(6508):739–746.

91. Kim C, Sano Y, Todorova K, Carlson BA, Arpa L, Celada A, Lawrence T, Otsu K, Brissette JL, Arthur JSC et al: The kinase p38α serves cell type–specific inflammatory functions in skin injury and coordinates pro– and anti-inflammatory gene expression. Nature immunology 2008, 9(9):1019–1027.

92. Noubade R, Wong K, Ota N, Rutz S, Eidenschenk C, Valdez PA, Ding J, Peng I, Sebrell A, Caplazi P et al: NRROS negatively regulates reactive oxygen species during host defence and autoimmunity. Nature 2014, 509(7499):235–239.

93. Ma W, Qin Y, Chapuy B, Lu C: LRRC33 is a novel binding and potential regulating protein of TGF-β1 function in human acute myeloid leukemia cells. PLOS ONE 2019, 14(10):e0213482.

94. Sehgal A, Irvine KM, Hume DA: Functions of macrophage colony-stimulating factor (CSF1) in development, homeostasis, and tissue repair. Seminars in Immunology 2021, 54:101509.

95. Larsson A, Carlsson L, Gordh T, Lind A-L, Thulin M, Kamali-Moghaddam M: The effects of age and gender on plasma levels of 63 cytokines. Journal of Immunological Methods 2015, 425:58–61.

96. Dinarello CA, Novick D, Kim S, Kaplanski G: Interleukin-18 and IL-18 Binding Protein. Frontiers in Immunology 2013, 4.

97. Rea IM, Gibson DS, McGilligan V, McNerlan SE, Alexander HD, Ross OA: Age and Age-Related Diseases: Role of Inflammation Triggers and Cytokines. Frontiers in Immunology 2018, 9.

98. Maeda M, Hasegawa H, Hyodo T, Ito S, Asano E, Yuang H, Funasaka K, Shimokata K, Hasegawa Y, Hamaguchi M et al: ARHGAP18, a GTPase-activating protein for RhoA, controls cell shape, spreading, and motility. Molecular Biology of the Cell 2011, 22(20):3840–3852.

99. Kapoor N, Gupta R, Menon ST, Folta-Stogniew E, Raleigh DP, Sakmar TP: Nucleobindin 1 Is a Calcium-regulated Guanine Nucleotide Dissociation Inhibitor of Gαi1. Journal of Biological Chemistry 2010, 285(41):31647–31660.

100. Su M-Y, Fromm SA, Remis J, Toso DB, Hurley JH: Structural basis for the ARF GAP activity and specificity of the C9orf72 complex. Nature Communications 2021, 12(1):3786.

101. Kometani K, Ishida D, Hattori M, Minato N: Rap1 and SPA-1 in hematologic malignancy. Trends in Molecular Medicine 2004, 10(8):401–408.

102. Krugmann S, Anderson KE, Ridley SH, Risso N, McGregor A, Coadwell J, Davidson K, Eguinoa A, Ellson CD, Lipp P et al: Identification of ARAP3, a Novel PI3K Effector Regulating Both Arf and Rho GTPases, by Selective Capture on Phosphoinositide Affinity Matrices. Molecular Cell 2002, 9(1):95–108.

103. Attwooll C, Denchi EL, Helin K: The E2F family: specific functions and overlapping interests. The EMBO journal 2004, 23(24):4709–4716.

104. Pitale PM, Gorbatyuk O, Gorbatyuk M: Neurodegeneration: Keeping ATF4 on a Tight Leash. Frontiers in Cellular Neuroscience 2017, 11.

105. Flury A, Aljayousi L, Park H-J, Khakpour M, Mechler J, Aziz S, McGrath JD, Deme P, Sandberg C, González Ibáñez F et al: A neurodegenerative cellular stress response linked to dark microglia and toxic lipid secretion. Neuron 2025, 113(4):554–571.e514.

106. Li X, Li Y, Jin Y, Zhang Y, Wu J, Xu Z, Huang Y, Cai L, Gao S, Liu T et al: Transcriptional and epigenetic decoding of the microglial aging process. Nature Aging 2023, 3(10):1288–1311.

107. Tan Q, Liang N, Zhang X, Li J: Dynamic Aging: Channeled Through Microenvironment. Frontiers in Physiology 2021, 12.

108. Sidler C, Wóycicki R, Ilnytskyy Y, Metz G, Kovalchuk I, Kovalchuk O: Immunosenescence is associated with altered gene expression and epigenetic regulation in primary and secondary immune organs. Frontiers in Genetics 2013, 4.

109. Franceschi C, Garagnani P, Parini P, Giuliani C, Santoro A: Inflammaging: a new immune–metabolic viewpoint for age-related diseases. Nature Reviews Endocrinology 2018, 14(10):576–590.

110. Jiao T, Chen Y, Sun H, Yang L: Targeting ferroptosis as a potential prevention and treatment strategy for aging-related diseases. Pharmacological research 2024, 208:107370.

111. Yang Y, Wang Y, Guo L, Gao W, Tang T-L, Yan M: Interaction between macrophages and ferroptosis. Cell Death & Disease 2022, 13(4):355.

112. Scott, Kathryn, Michael, Skouta R, Eleina, Caroline, Darpan, Andras, Alexandra, Wan et al: Ferroptosis: An Iron-Dependent Form of Nonapoptotic Cell Death. Cell 2012, 149(5):1060–1072.

113. Zeidan RS, Han SM, Leeuwenburgh C, Xiao R: Iron homeostasis and organismal aging. Ageing research reviews 2021, 72:101510.

114. Sato T, Shapiro JS, Chang H-C, Miller RA, Ardehali H: Aging is associated with increased brain iron through cortex-derived hepcidin expression. ELife 2022, 11.

115. Ward RJ, Zucca FA, Duyn JH, Crichton RR, Zecca L: The role of iron in brain ageing and neurodegenerative disorders. The Lancet neurology 2014, 13(10):1045–1060.

116. Handa P, Thomas S, Morgan-Stevenson V, Maliken BD, Gochanour E, Boukhar S, Yeh MM, Kowdley KV: Iron alters macrophage polarization status and leads to steatohepatitis and fibrogenesis. Journal of leukocyte biology 2019, 105(5):1015–1026.

117. Zhou Y, Que KT, Zhang Z, Yi ZJ, Zhao PX, You Y, Gong JP, Liu ZJ: Iron overloaded polarizes macrophage to proinflammation phenotype through ROS /acetyl-p53 pathway. Cancer medicine 2018, 7(8):4012–4022.

118. Inestrosa NC, Tapia-Rojas C, Lindsay CB, Zolezzi JM: Wnt Signaling Pathway Dysregulation in the Aging Brain: Lessons From the Octodon degus. Frontiers in Cell and Developmental Biology 2020, 8.

119. García-Velázquez L, Arias C: The emerging role of Wnt signaling dysregulation in the understanding and modification of age-associated diseases. Ageing Research Reviews 2017, 37:135–145.

120. Yu X, Wang Y, Song Y, Gao X, Deng H: AP-1 is a regulatory transcription factor of inflammaging in the murine kidney and liver. Aging Cell 2023, 22(7):e13858.

121. Karakaslar EO, Katiyar N, Hasham M, Youn A, Sharma S, Chung CH, Marches R, Korstanje R, Banchereau J, Ucar D: Transcriptional activation of Jun and Fos members of the AP-1 complex is a conserved signature of immune aging that contributes to inflammaging. Aging Cell 2023, 22(4).

122. Zimmerman SM, Kim SK: The GATA transcription factor/MTA-1 homolog egr-1 promotes longevity and stress resistance in Caenorhabditis elegans. Aging Cell 2014, 13(2):329–339.

123. Hecker D, Lauber M, Behjati Ardakani F, Ashrafiyan S, Manz Q, Kersting J, Hoffmann M, Schulz MH, List M: Computational tools for inferring transcription factor activity. Proteomics 2023, 23(23-24).

124. Palladino I, Salani F, Ciaramella A, Rubino IA, Caltagirone C, Fagioli S, Spalletta G, Bossù P: Elevated levels of circulating IL-18BP and perturbed regulation of IL-18 in schizophrenia. Journal of Neuroinflammation 2012, 9(1):206.

125. Davies ML, Biryukov SS, Rill NO, Klimko CP, Hunter M, Dankmeyer JL, Miller JA, Shoe JL, Mlynek KD, Talyansky Y et al: Sex differences in immune protection in mice conferred by heterologous vaccines for pneumonic plague. Frontiers in Immunology 2024, 15.

126. Pittet J-F, Hu PJ, Honavar J, Brandon AP, Evans CA, Muthalaly R, Ding Q, Wagener BM: Estrogen Alleviates Sex-Dependent Differences in Lung Bacterial Clearance and Mortality Secondary to Bacterial Pneumonia after Traumatic Brain Injury. Journal of Neurotrauma 2021, 38(8):989–999.

127. Gubbels Bupp MR, Potluri T, Fink AL, Klein SL: The Confluence of Sex Hormones and Aging on Immunity. Frontiers in Immunology 2018, 9.

128. Kessels S, Trippaers C, Mertens M, Hamad I, Rombaut B, Janssen A, Ramanathan K, Duwé S, Gharghani AM, Theuwis R et al: Cytoskeletal control in adult microglia is essential to restore neurodevelopmental synaptic and cognitive deficits. Science Advances 2025, 11(35).

129. Elmore MRP, Hohsfield LA, Kramár EA, Soreq L, Lee RJ, Pham ST, Najafi AR, Spangenberg EE, Wood MA, West BL et al: Replacement of microglia in the aged brain reverses cognitive, synaptic, and neuronal deficits in mice. Aging Cell 2018, 17(6):e12832.

130. Socodato R, Relvas JB: A cytoskeleton symphony: Actin and microtubules in microglia dynamics and aging. Progress in Neurobiology 2024, 234:102586.

131. Ghannam A, Hahn V, Fan J, Tasevski S, Moughni S, Li G, Zhang Z: Sex-specific and cell-specific regulation of ER stress and neuroinflammation after traumatic brain injury in juvenile mice. Experimental neurology 2024, 377:114806.

132. Ching T, Huang S, Garmire LX: Power analysis and sample size estimation for RNA-Seq differential expression. RNA 2014, 20(11):1684–1696.

133. Singh PP, Benayoun BA: Considerations for reproducible omics in aging research. Nature Aging 2023, 3(8):921–930.

134. Liao Y, Smyth GK, Shi W: featureCounts: an efficient general purpose program for assigning sequence reads to genomic features. Bioinformatics 2014, 30(7):923–930.

135. Leek JT, Storey JD: Capturing Heterogeneity in Gene Expression Studies by Surrogate Variable Analysis. PLoS Genetics 2007, 3(9):e161.

136. Ritchie ME, Phipson B, Wu DI, Hu Y, Law CW, Shi W, Smyth GK: limma powers differential expression analyses for RNA-sequencing and microarray studies. Nucleic acids research 2015, 43(7):e47–e47.

137. Love MI, Huber W, Anders S: Moderated estimation of fold change and dispersion for RNA-seq data with DESeq2. Genome biology 2014, 15(12):550.

138. Hao Y, Stuart T, Kowalski MH, Choudhary S, Hoffman P, Hartman A, Srivastava A, Molla G, Madad S, Fernandez-Granda C et al: Dictionary learning for integrative, multimodal and scalable single-cell analysis. Nature Biotechnology 2024, 42(2):293–304.

139. Yu G, Wang L-G, Han Y, He Q-Y: clusterProfiler: an R Package for Comparing Biological Themes Among Gene Clusters. OMICS: A Journal of Integrative Biology 2012, 16(5):284–287.

140. Wickham H: Data analysis. In: ggplot2: elegant graphics for data analysis. Springer; 2016: 189–201.

141. Milacic M, Beavers D, Conley P, Gong C, Gillespie M, Griss J, Haw R, Jassal B, Matthews L, May B et al: The Reactome Pathway Knowledgebase 2024. Nucleic Acids Research 2024, 52(D1):D672–D678.

142. Dolgalev I: MSigDB Gene Sets for Multiple Organisms in a Tidy Data Format. 2025.

143. Müller-Dott S, Tsirvouli E, Vazquez M, Ricardo, Badia-I-Mompel P, Fallegger R, Türei D, Lægreid A, Saez-Rodriguez J: Expanding the coverage of regulons from high-confidence prior knowledge for accurate estimation of transcription factor activities. Nucleic Acids Research 2023, 51(20):10934–10949.

