## Supplementary figures and images for "Shared and niche-specific transcriptional signatures of macrophage aging revealed by a cross-tissue meta-analysis"

### Supplemental Figures 1-8

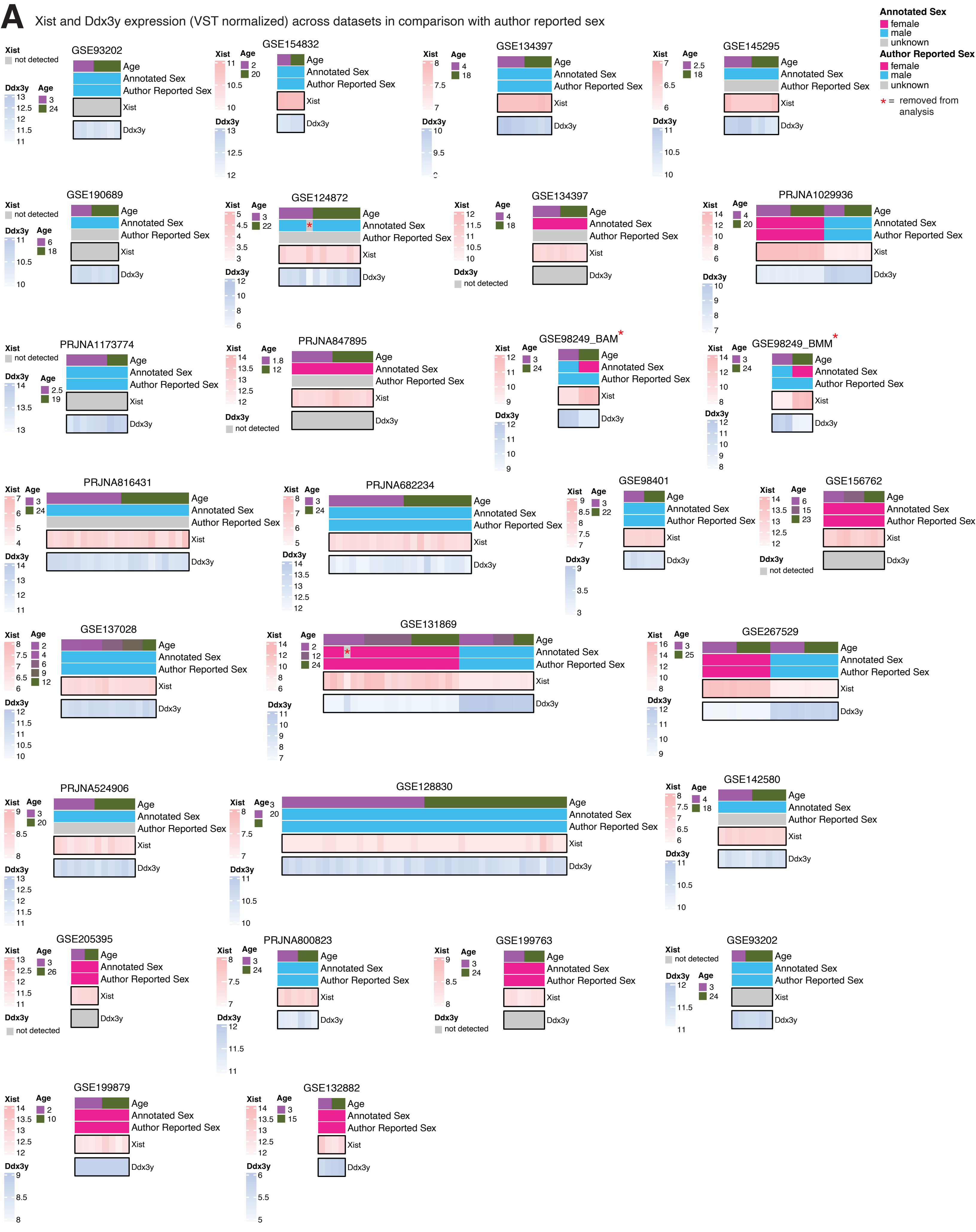

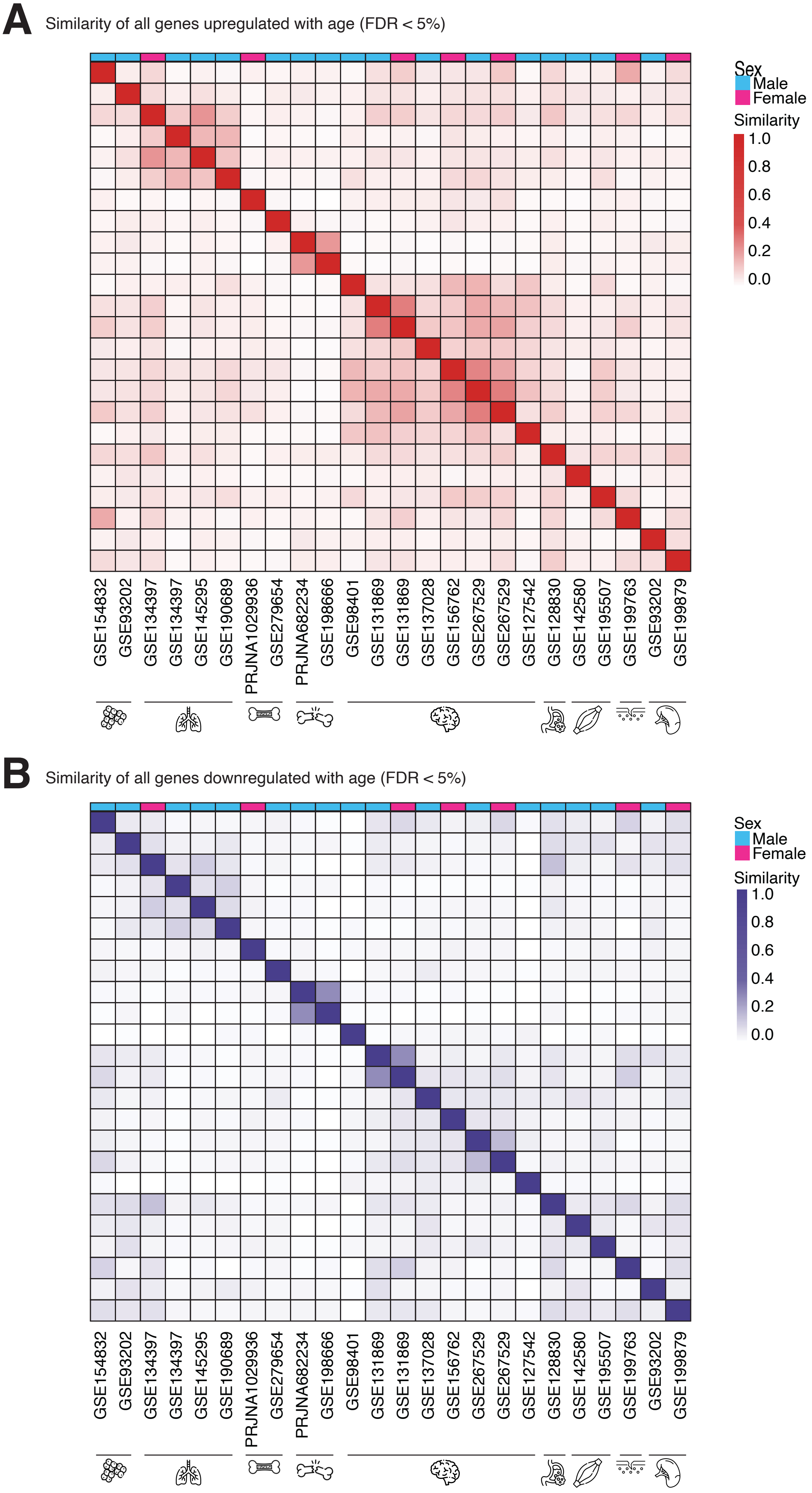

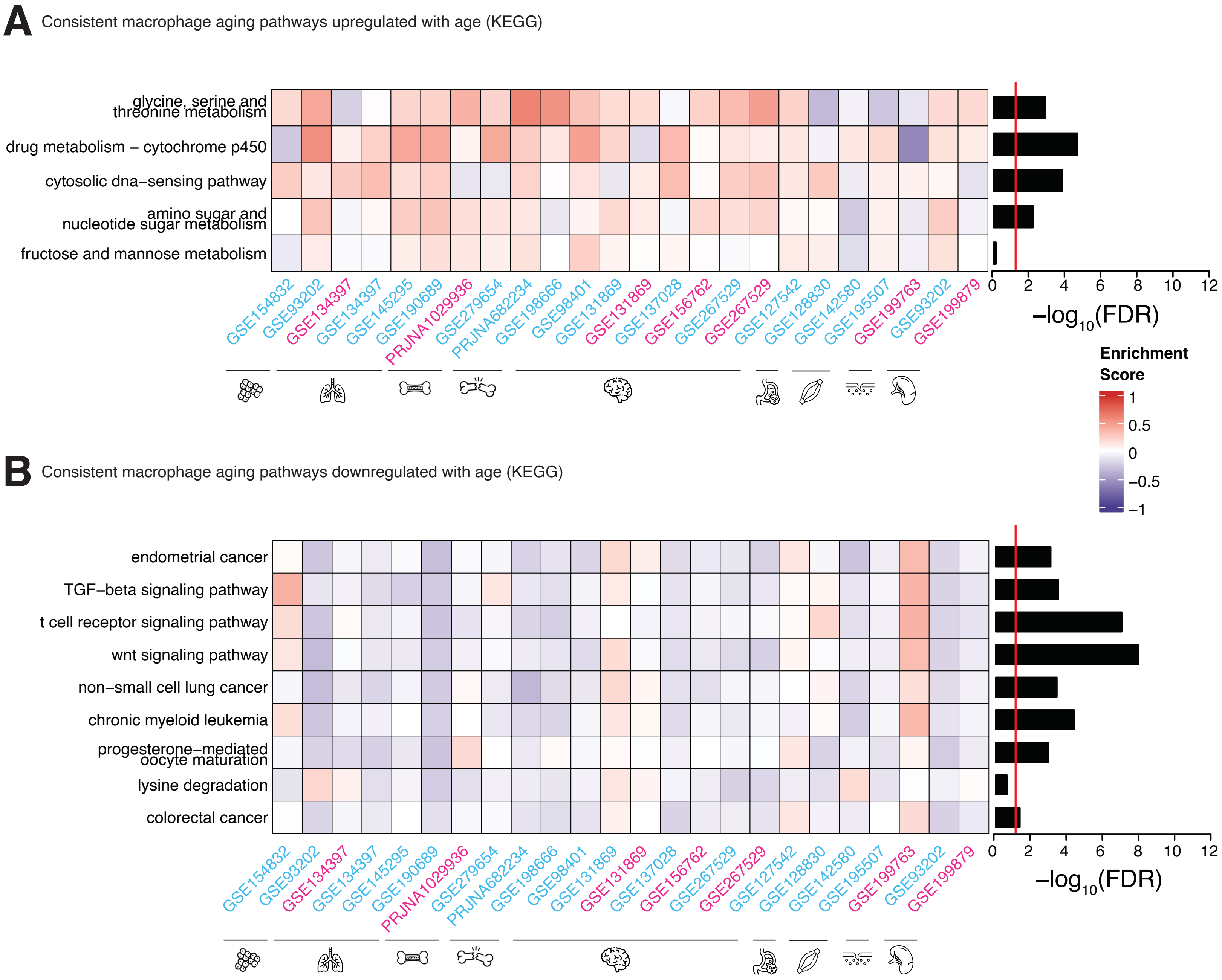

**A** Macrophage aging transcription factor-pathway network scores (KEGG)

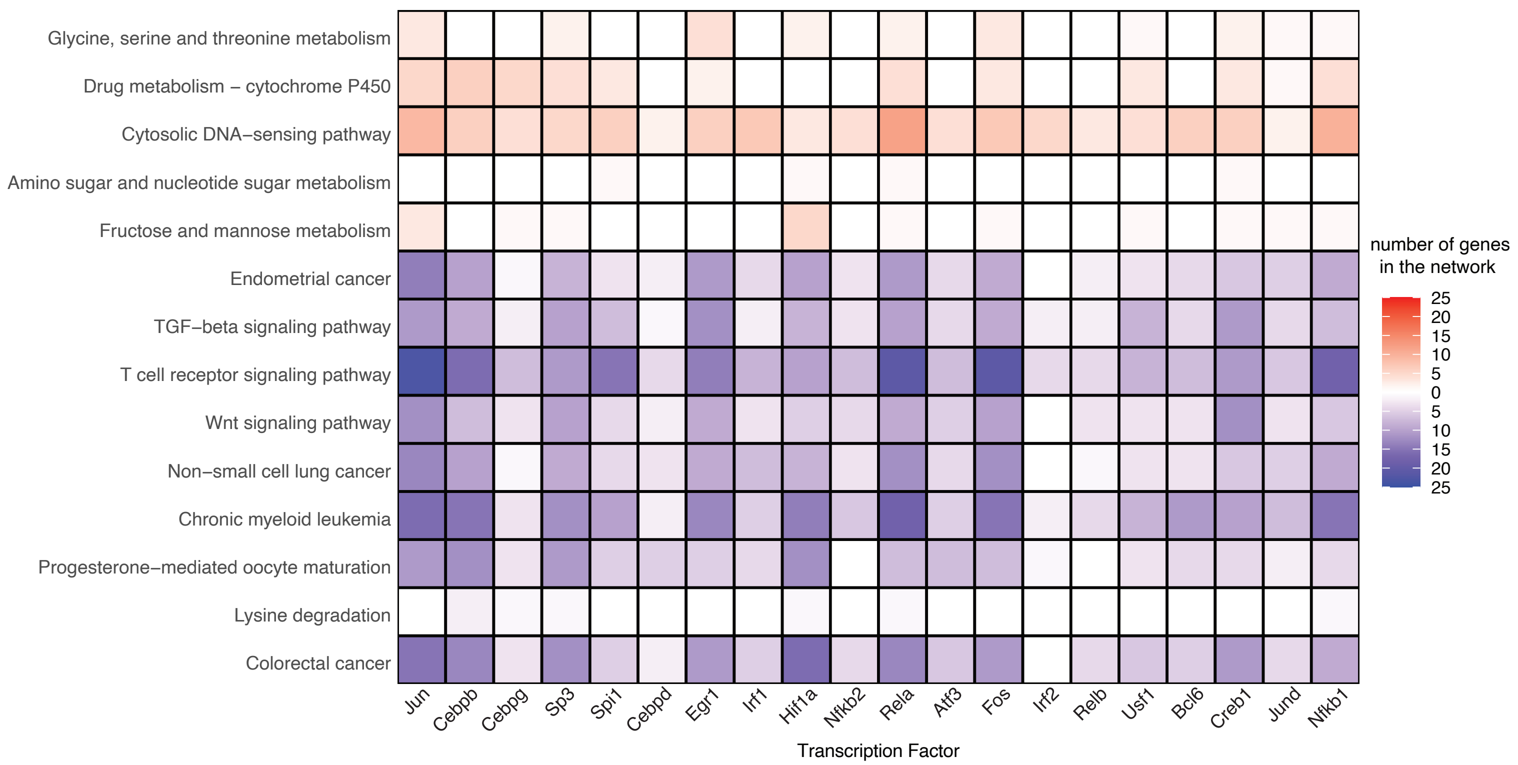

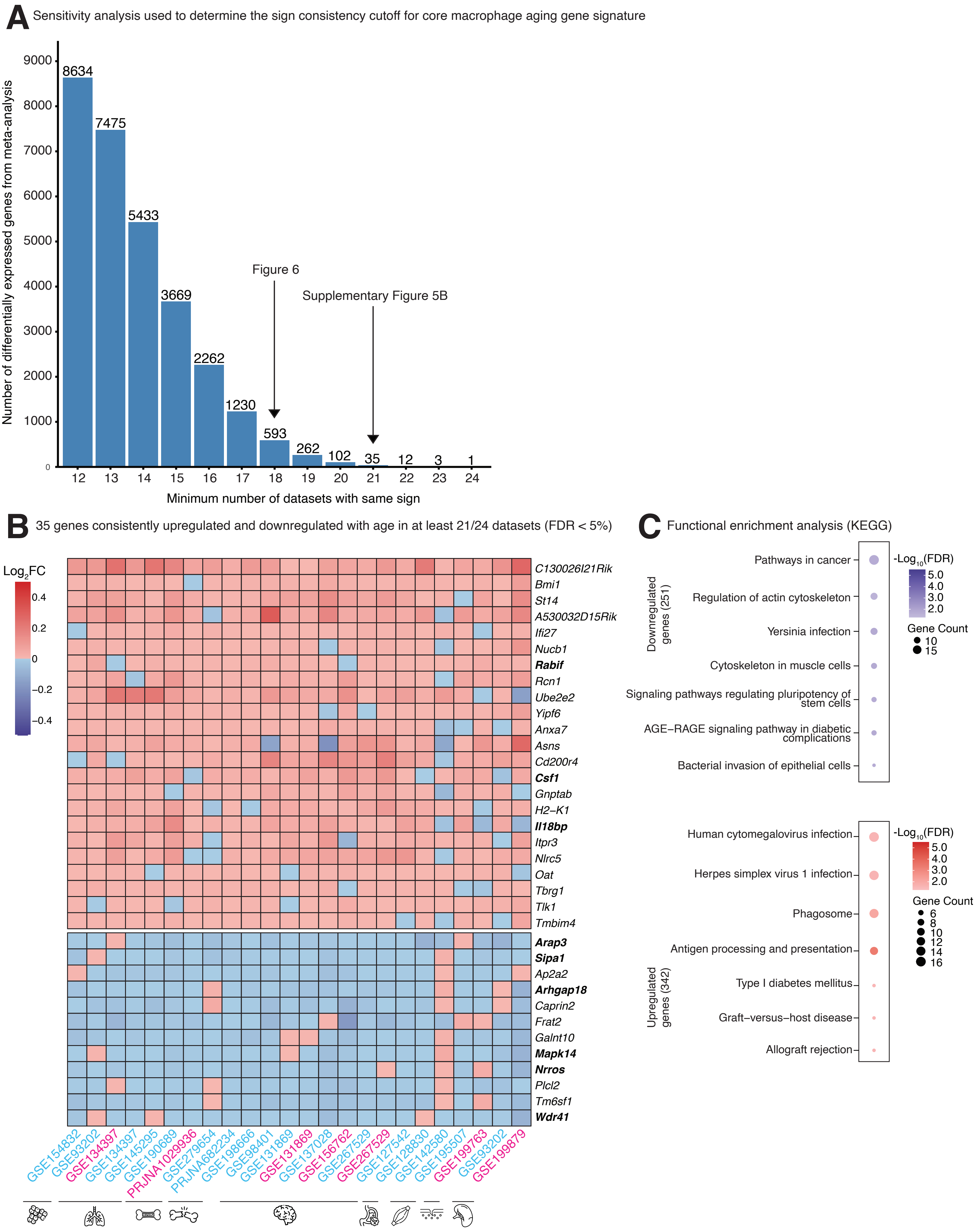

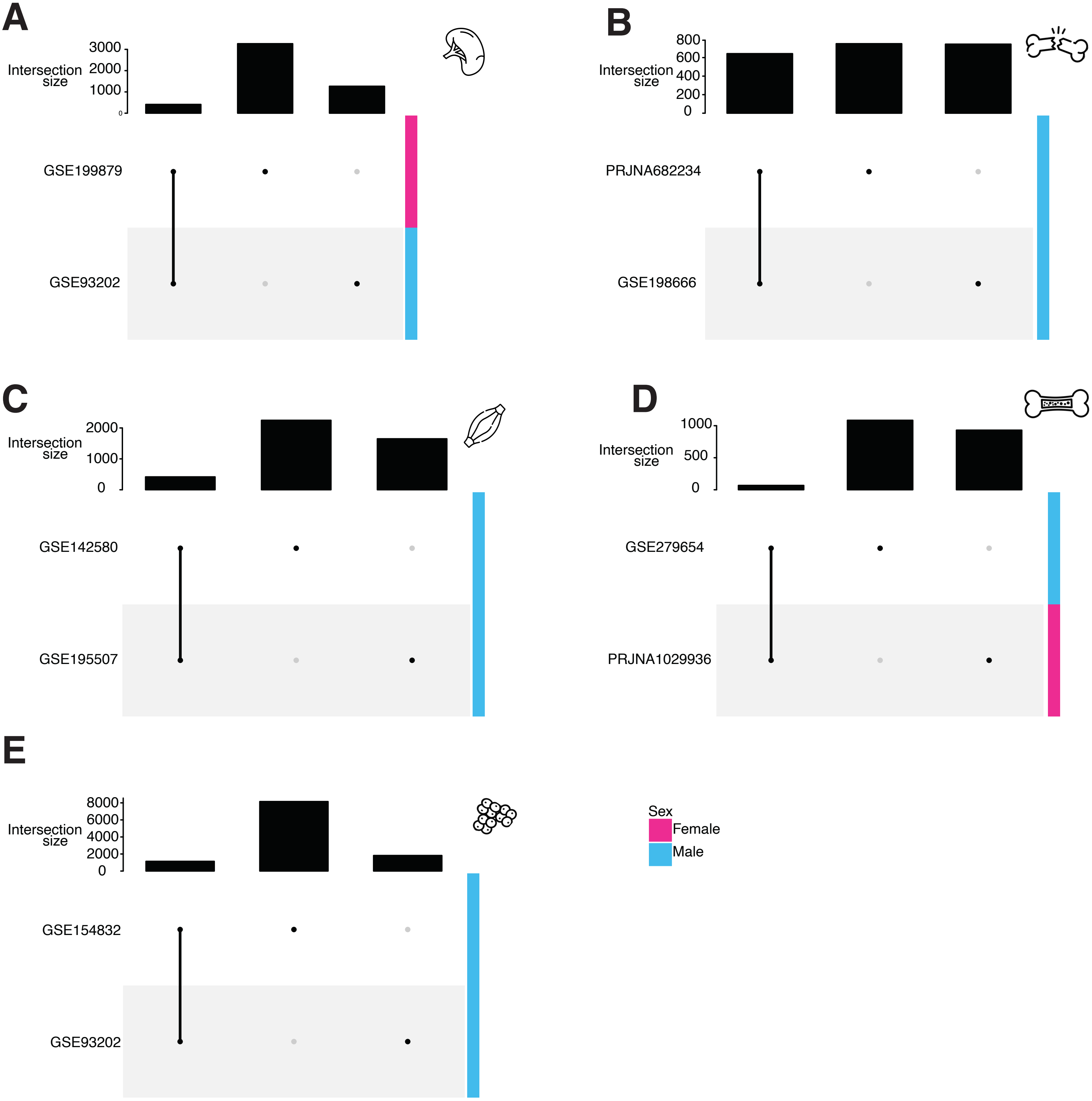

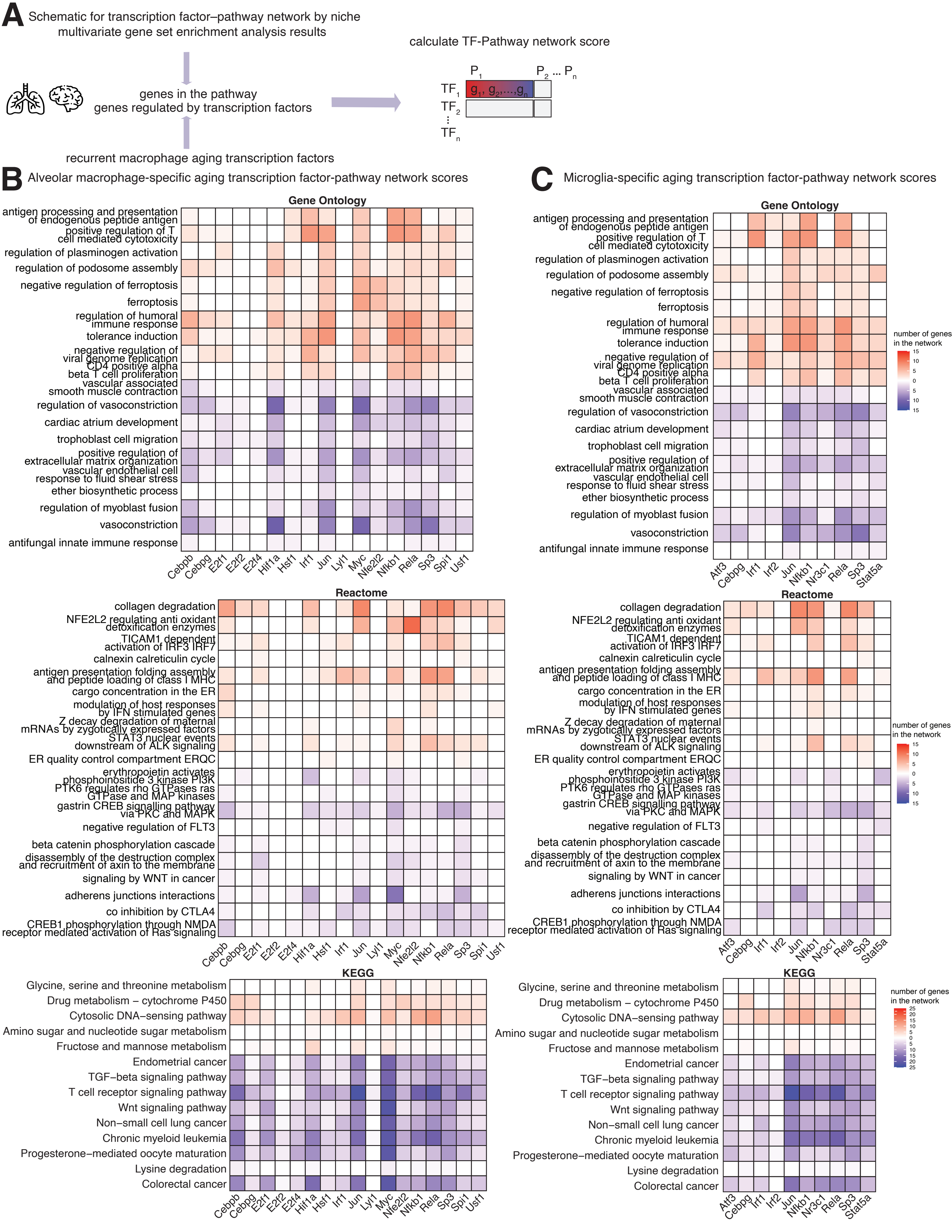

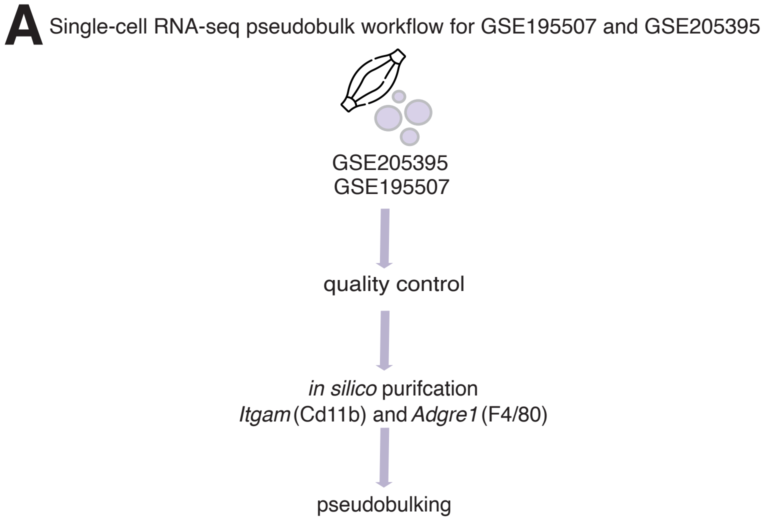
